# Astrocytes connect specific brain regions through plastic gap junctional networks

**DOI:** 10.1101/2025.07.18.665573

**Authors:** Melissa L Cooper, Maria Clara Selles, Michael Cammer, Holly K Gildea, Joseph Sall, Katelyn E Chiurri, Aiman S Saab, Shane A Liddelow, Moses V Chao

**Author notes:** Jointly supervised this work (Shane A Liddelow, Moses V Chao).

## Abstract

Traditionally, neuronal axons have been considered the primary mediators of functional connectivity among brain regions. However, the role of astrocyte-mediated communication has been largely underappreciated. While astrocytes communicate with one another through gap junctions, the extent and specificity of this communication remain poorly understood. Astrocyte gap junctions are necessary for memory formation^1,2^, synaptic plasticity^3-5^, coordination of neuronal signaling^6^, and closing the visual and motor critical periods^7,8^. These findings indicate that this form of communication is essential for proper central nervous system development and function. Despite their significance, studying astrocyte gap junctional networks has been challenging. Current methods like slice electrophysiology disrupt network connectivity and introduce artifacts due to tissue damage. To overcome these limitations, we developed a vector-based approach that labels molecules as they are fluxed by astrocyte gap junctions in awake, behaving animals. We then used whole-brain tissue clearing^9,10^ to image these intact, three-dimensional astrocyte networks. We show that multiple astrocyte networks traverse the mouse brain. These networks selectively connect specific regions, rather than diffusing indiscriminately, and vary in size and organization. We observe local networks are confined to single brain regions and long-range networks robustly interconnecting multiple regions across hemispheres, often exhibiting patterns distinct from known neuronal networks. Further, we demonstrate that astrocyte networks undergo structural reorganization in adult brain following sensory deprivation. These discoveries reveal a previously unrecognized mode of communication between distant brain regions, mediated by plastic networks of gap junction-coupled astrocytes.

## INTRODUCTION

Astrocyte intercellular communication is critical to proper central nervous system (CNS) function. This communication occurs through gap junctions, membrane channels connecting the cytoplasm of neighboring cells that enable them to redistribute resources and share biochemical signals. Studies using astrocyte gap junction knockout mice have shown their necessity for memory formation^1,2^, synaptic plasticity^3-5^, coordination of neuronal signaling^6^, and closing the visual and motor critical periods^7,8^. In disease, networks of gap junction linked astrocytes redistribute metabolic resources across the CNS to protect degenerating neurons^11,12^. Despite these insights, our understanding of the spatial architecture and functional topology of astrocyte networks remains limited. Existing methods, such as dye diffusion in acute slices or reporter activation in injury models, are inherently constrained to local environments and often disrupt native connectivity. As a result, it remains unknown whether astrocytes form a continuous brain-wide syncytium or operate through discrete, region-specific subnetworks. It is also unclear whether their anatomical connectivity aligns with neuronal networks or establishes an independent framework for long-range, non-neuronal signaling.

## RESULTS

### Astrocyte gap-junctional network tracing

To address this knowledge gap, we developed a vector-based approach to express a fusion protein comprised of connexin 43 (Cx43), the main gap junction protein used by astrocytes, and TurboID, a rapid and promiscuous biotinylating enzyme^13-15^, under the shortened *Gfap* promoter^16-18^ (AAV5-GfaABC1D-Cx43:TID:HA; Figure 1a). When this fusion protein incorporates into a connexon as any one of its 6 constituent connexins (Figure 1b), molecules that flux through the infected astrocyte ‘s gap junctions are rapidly tagged with small, inert biotin (Figure 1c). This enables us to detect the infected astrocyte population (via the HA tag on the fusion protein), the in-network astrocytes (through staining biotinylated moieties with streptavidin), and cells that are not in-network with the infected population (Figure 1d). Because the mouse CNS has minimal-to-no native biotin^13-15^, streptavidin staining provides little background. Further, each biotin can only bind a single streptavidin, eliminating variability introduced through antibody multiplexing and making images more quantitative.

**Figure 1.**
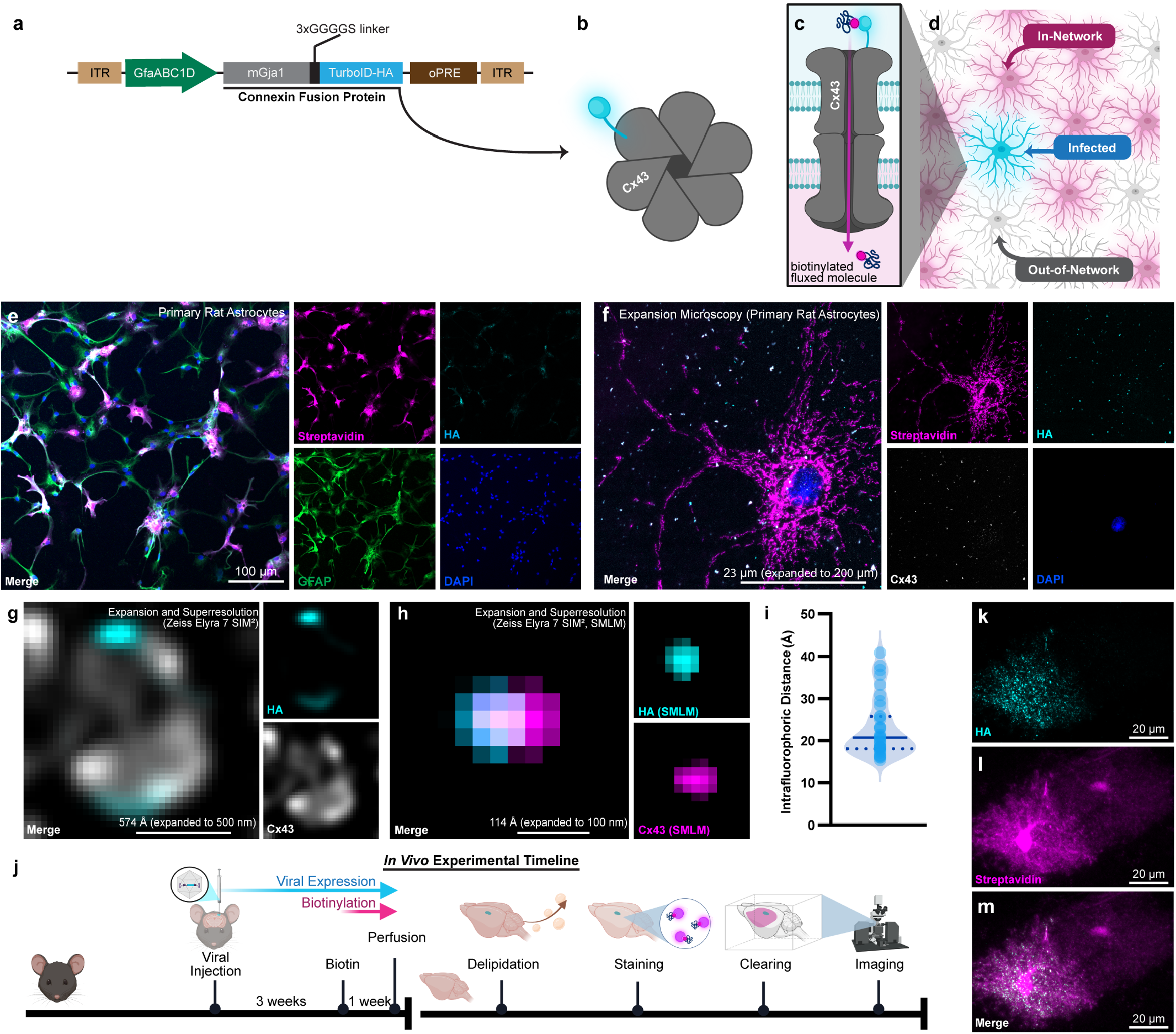
Visualizing astrocyte gap junctional communication using the astrocyte network tracer. **a**. Diagram of the astrocyte network tracer construct. **b**. Cx43 connexons contain 6 connexins, only one of which needs to be the fusion protein for TurboID to reside on the gap junction. **c**. Infected astrocytes biotinylate molecules that flux through Cx43 gap junctions into adjacent, uninfected cells. **d**. This volume-fills in-network astrocytes with biotinylated molecules such that infected (HA+, streptavidin+), in-network (HA-, streptavidin+), and out-of-network (HA-, streptavidin-) cells can be visualized. **e**. Infection of primary immunopanned rat astrocytes (GFAP, green) resulted in Cx43-TID-HA expression (cyan) in 10% of cells, forming a biotin-labeled network containing 80% of cells (streptavidin, magenta) (see Supplemental Figures 1, 2). f. Primary rat astrocytes were expanded 8.7x to visualize individual Cx43 gap junctions (grey) and incorporated fusion protein (HA, cyan). **g**. Superresolution microscopy on expanded astrocytes^20^ enabled single-molecule imaging of the fusion protein incorporated into astrocyte gap junctions expressed in immunopanned primary rat astrocytes. Polyclonal Cx43 antibodies conjugated directly to fluorophores reveal gap junction structure (grey) and the HA tag (cyan) shows TurboID within the gap junction vestibule. **h**. Single-molecule localization microscopy (SMLM) of HA (cyan) and Cx43 (magenta). **i**. Quantification of the intrafluorophoric distance from HA to the nearest Cx43. Median (20.73 Å) indicated with the solid blue line; dotted lines denote the 25th (18.09 Å) and 75th (25.74 Å) percentiles. **j**. In vivo experimental timeline. 3 weeks following viral injection, mice received biotin-supplemented drinking water for 1 week, after which they were perfused. Brains were then delipidated, stained with streptavidin to reveal biotinylated molecules, cleared, and imaged via light sheet microscopy. **k**. Light sheet micrograph of HA-tag (cyan) shows gap junctions containing fusion protein in an infected cell within a cleared brain. l. Streptavidin (magenta) labeling of biotinylated molecules. **m**. Merged image showing a two-cell example of an infected (left) and an uninfected in-network (right) cell.

To validate the construct, we first infected cultures of primary immunopanned serum-free rat astrocytes (Figure 1e; purity confirmation in Supplemental Figure 1a). When ∼10% of cells expressed Cx43-TID, over 80% of cells in the plate were identifiably in-network (Figure 1e; quantification and controls in Supplemental Figure 2). Streptavidin staining beyond the infected astrocyte population was confirmed to be gap junction-dependent in primary astrocyte cultures from inducible astrocyte gap junction knockout mice (Supplemental Figures 1, 3).

To confirm that TurboID resides within the gap junction vestibule, where it would be proximal to molecules staged for gap junction flux^19^, we used a combination of expansion microscopy^20^ (Figure 1f) and superresolution imaging (Zeiss Elyra 7 SIM2). This approach allowed us to visualize the fusion protein incorporated within a gap junction in expanded primary cultured rat astrocytes^21^ (Figure 1g). It also enabled us to measure the distance between the Cx43 vestibule and the HA tag on the N terminus of TurboID (Figure 1h,i). The 20.73 Å median distance indicates that the HA tag lies immediately outside of the ∼15 Å gap junction vestibule^19^. While the 3D structure of TurboID has not been solved, the labeling radius of biotin ligases throughout the TurboID family of enzymes has been estimated at ∼10-30 nm (100-300 Å)^13,22,23^, well within all measured intrafluorophoric distances.

Following our in vitro validation of the fusion protein ‘s structure and function, we moved to in vivo experiments (Figure 1j). For each in vivo experiment, mice were unilaterally injected with AAV5-GfaABC1D-Cx43:TID:HA (henceforth referred to as ‘astrocyte network tracer ‘) in a single defined brain region. Three weeks later, biotin was administered via drinking water for one week. On day 28, mice were perfused and brains carefully dissected. Brains were then delipidated, stained, cleared, and imaged using light sheet microscopy. In cleared brains, infected cells expressed Cx43-TID-HA in a punctate pattern on the cell membrane, characteristic of gap junctions (Figure 1k). Streptavidin staining revealed an adjacent in-network, uninfected cell (Figure 1l); while this cell was negative for Cx43-TID-HA, it contained biotinylated molecules that had passed through the infected astrocyte ‘s gap junctions (Figure 1m).

### Astrocyte networks vary regionally

To visualize in vivo astrocyte networks, assess their reproducibility across mice, and examine regional differences in connectivity, we injected male, 3-month-old C57BL/6 mice in one of 3 brain regions: motor cortex (Figure 2, top row), hypothalamus (Figure 2, middle row), or prefrontal cortex (Figure 2, bottom row). Dorsal, sagittal, and oblique views of the light sheet datasets illustrate distinct network morphologies across regions and reveal the specificity of astrocytic connectivity. Infected cells displayed the canonical stellate morphology of astrocytes as well as the expected punctate HA-tag staining pattern (Supplemental Figure 4). We found consistent patterns of regional connectivity among animals injected in the same area (Supplemental Figures 5, 6, 7), suggesting that astrocyte networks are conserved across mice.

**Figure 2.**
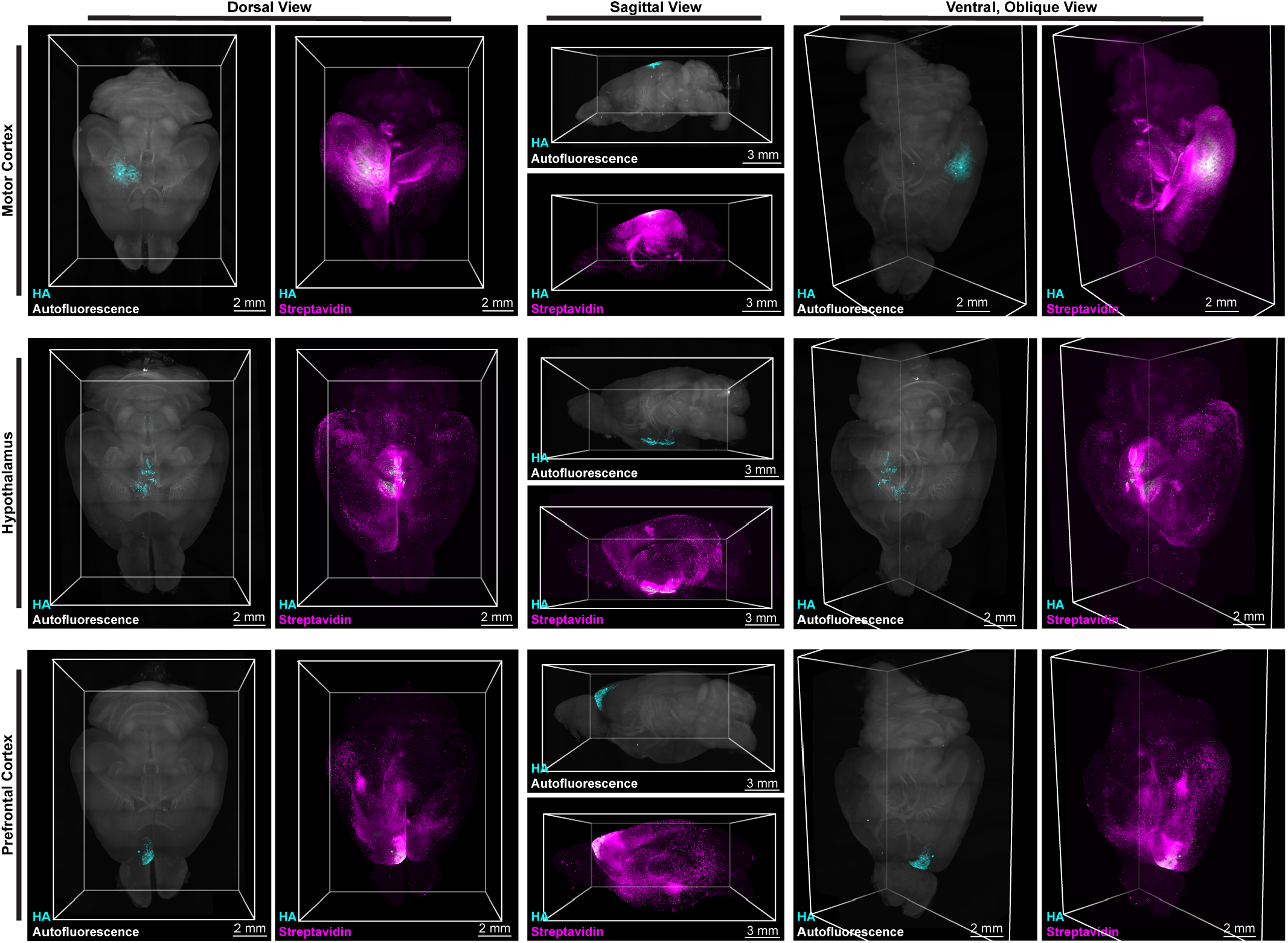
Multiple astrocyte networks varying in size and organization traverse the mouse brain. Three-dimensional renderings of light sheet-imaged brains infected in motor cortex (top row), hypothalamus (middle row), or prefrontal cortex (bottom row). Each image is presented in two views: the first shows the infected region (HA-tag, cyan) inside of the autofluorescence capture (grey) for context; the second shows the same infected region within the streptavidin-stained astrocyte network originating from that brain region (magenta). Dorsal (left column), sagittal (middle column), and oblique (right column) views of the same light sheet datasets show network morphology at different angles.

Virtual coronal sections of these brains, mapped^24^ to the Allen Brain Atlas, further illustrate the specificity and complexity of astrocyte networks (Figure 3). Regions with robust connectivity are often directly adjacent to areas that lack any detectable connections. Within a single astrocyte network, different brain regions can display distinct patterns of network-linked cells (Supplemental Figure 8). While some regions remain largely unilaterally connected (Supplemental Figure 8b,e), others show robust bilateral connectivity (Supplemental Figure 8c). In rare instances, specific neuronal populations contain biotinylated molecules. When present, this labeling typically occurs at the terminus of an interconnected chain of astrocytes and is restricted to neurons within a localized area (Supplemental Figure 8k). Most cells (and in many networks, all cells) that contain biotinylated molecules exhibit the characteristic stellate morphology of astrocytes.

**Figure 3.**
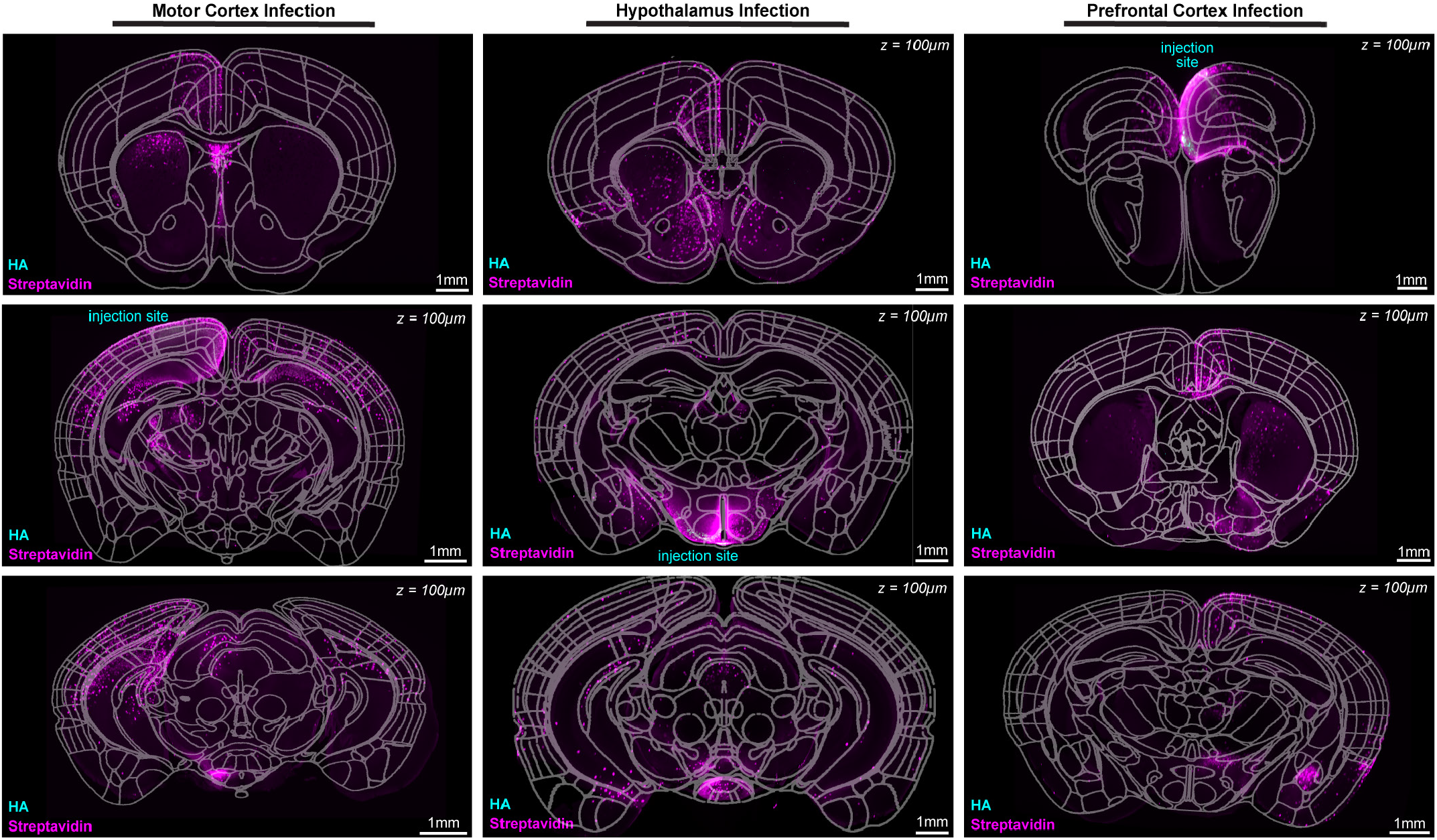
Astrocyte networks link specific brain regions. Virtual coronal sections of light sheet imaged brains infected in motor cortex (left column), hypothalamus (middle column), or prefrontal cortex (right column). Sections from each brain are arranged from rostral to caudal (top to bottom images) and were selected as representative examples of robust astrocyte network connections. Brain regions are outlined using an overlay of the Allen Brain Atlas (fitted via the FIJI ABBA plugin^24^; 3D mouse Allen Brain Atlas, mouse.brain-map.org and atlas.brain-map.org).

Astrocyte networks can link multiple astrocyte subtypes. In bilateral networks, chains of interconnected astrocytes spanning white matter tracts such as corpus collosum are clearly visible (Supplemental Figure 9). Rather than filling the entire corpus callosum, these chains of astrocytes appear to follow specific neuronal axons. Networks can also include both parenchymal astrocytes and glia limitans superficialis astrocytes^25^ located at the brain ‘s surface (Supplemental Figure 10). These two astrocytic populations contact one another, although the glia limitans superficialis astrocytes are not necessarily directly coupled to one another.

### Gap junctions link distant astrocytes

To examine whether non-gap junctional mechanisms could account for biotinylated molecules beyond the initially infected cells, we used tamoxifen-inducible, astrocyte-specific Cx30 and Cx43 double knockouts (*Slc1a3*(GLAST)^Cre-ERT2+/^*Gja1*(Cx43)^fl/fl^*Gjb6*(Cx30)^fl/fl^, referred to herein as cKO)^1^. Knockout of Cx43 alone is insufficient, as astrocytes also express connexin 30 (Cx30), albeit at lower levels^1^. We designed our vector to biotinylate molecules fluxed by Cx43 because it is the most permissive connexin^26^. However, we knocked out both Cx43 and Cx30 in this cKO to control for the possibility that a subset of small molecules might still pass through Cx30 in the absence of Cx43.

We first used immunohistochemistry (Figure 4, Supplemental Figure 3) and qPCR (Supplemental Figure 1) to confirm reduction of connexin expression in cKO mouse astrocytes. One week after tamoxifen gavage, immunoreactivity for Cx43 (Figure 4a,c,e) and Cx30 (Figure 4b,d,f) was diminished in cKO mice (right column) relative to littermate controls (left column). As expected for Cx43, a small degree of immunoreactivity was retained around vasculature in cKO mice (vasculature marked in Figure 4c-f using Lycopersicon Esculentum (Tomato) lectin (TL), red), reflecting pericyte localization. Pericytes express Cx43^27^ and are not Cre+ in this model. The persistence of connexins expression adjacent to vasculature, but not elsewhere, confirms that our gap junction knockout was specific to astrocytes. We quantified this by determining fluorescence intensity relative to the distance from the nearest blood vessel in two-dimensional sections. Immunoreactivity in cKO brains (n = 3 per group) was significantly reduced compared to littermate controls at all locations except for ∼3μm from vessel walls, where pericytes reside (Figure 4i).

**Figure 4.**
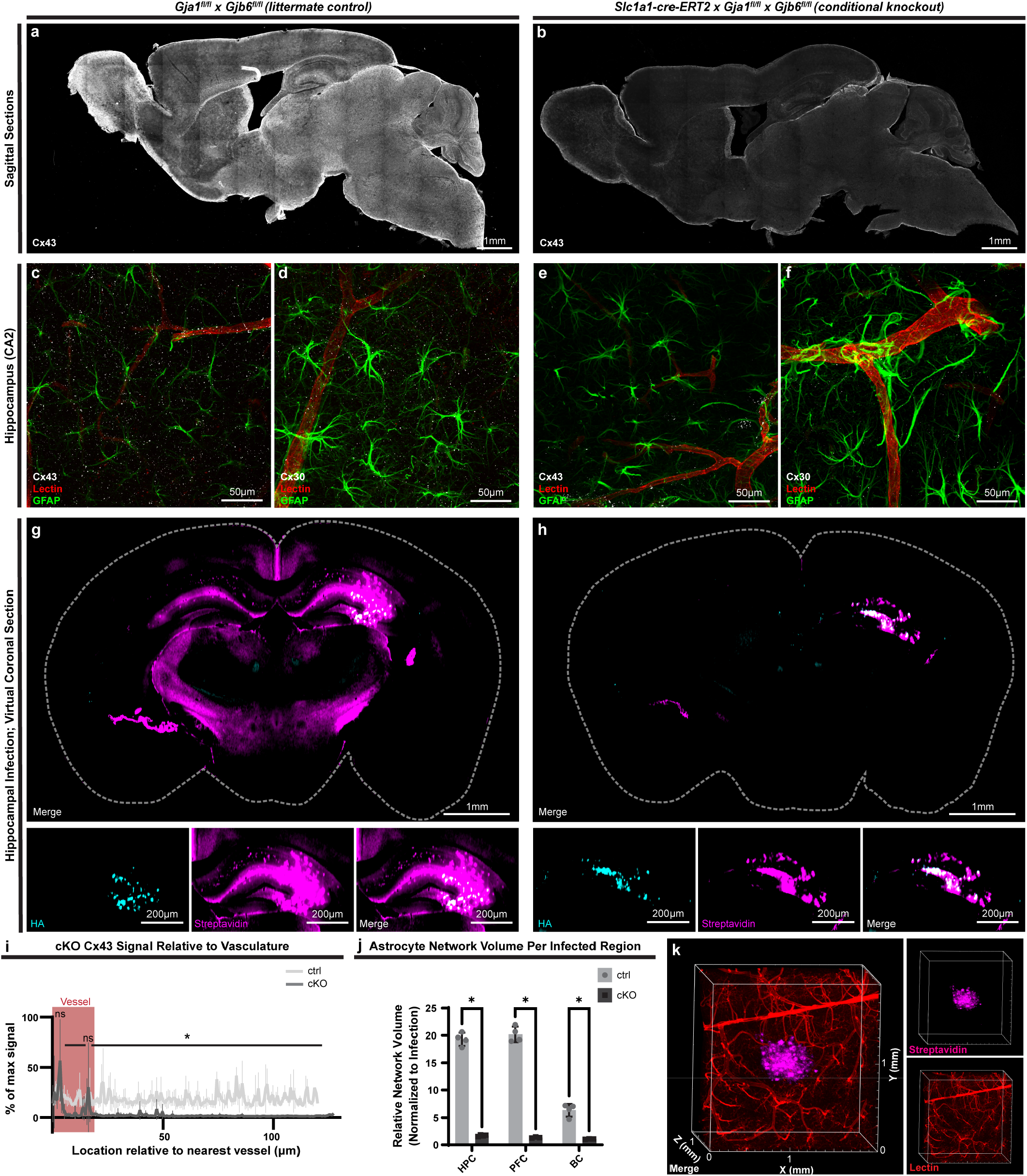
Non-gap-junctional mechanisms do not compensate for astrocyte gap junctional knockout. **a**. Sagittal section of *Cx43*^fl/fl^*Cx30*^fl/fl^ mouse brain, littermate to **b**. *Slc1a1*(GLAST)^Cre-ERT2^*Gja1*^fl/fl^*Gjb6*^fl/fl^ (cKO) mouse. Both mice were given tamoxifen, inducing robust Cx ablation in cKO mice; sections are stained for Cx43. **c-f**. Higher resolution images of hippocampi stained for GFAP (green, astrocytes), tomato lectin (red, vasculature), and either Cx43 or Cx30 (white) in control (c, d) or conditional knockout (e, f) mice. **g**. Control mouse injected in hippocampus expresses the astrocyte network tracer in CA2 (HA tag, cyan) and exhibits bilateral streptavidin labeling across hippocampus and hypothalamic regions. **h**. In conditional knockout mice, streptavidin labeling is restricted to infected astrocytes where Cx43 is reintroduced via the astrocyte network tracer. **i**. Quantification of Cx43 levels across vasculature and parenchyma. In cKO mice, Cx43 immunoreactivity is significantly diminished in all locations except at the periphery of blood vessels, where pericytes reside (paired two-sided t-tests across 180 locations relative to vasculature with Bonferroni correction; p < 0.001 in all significant locations; n = 3 mice per condition with 3 micrographs averaged to generate values for each mouse). **j**. Quantification of astrocyte network volume normalized to the volume of infected astrocytes. When astrocytes are infected in either hippocampus, prefrontal cortex (PFC), or barrel cortex, conditional knockout mouse networks are significantly reduced from those in control mice (unpaired two-sided t-tests; p < 0.001 in all regions; n = 4 mice per region per condition). **k**. cKO mice induced via tamoxifen gavage were injected with astrocyte network tracer per the experimental timeline in Figure 1j. Tomato Lectin was injected during perfusion to stain vasculature. Micrographs show the streptavidin-labeled astrocyte network (magenta) and vasculature (red). The network does not appear to follow or expand along blood vessels following astrocyte network loss.

We infected 4 cKO and 4 control mice per condition in either hippocampus (HPC), prefrontal cortex (PFC), or barrel cortex (BC). Astrocyte networks originating in each region were significantly smaller in cKO mice compared to controls (Figure 4j). Network volume within each brain region was consistent among animals in the same condition, even among networks of different sizes, such as the relatively small barrel cortex astrocyte network. This internal consistency parallels that observed in C57BL/6 mice injected in the motor cortex, prefrontal cortex, and hypothalamus (Supplemental Figures 5, 6, 7). Following knockout, network volume in each brain region was restricted to the infected astrocytes, which had Cx43 reintroduced via the astrocyte network tracer. These cells could form a network with one another, but not with neighboring uninfected astrocytes lacking Cx43 and Cx30.

We next examined whether pericyte-expressed Cx43 or other vasculature-related mechanisms could compensate for the loss of astrocyte gap junctions and support the distribution of biotinylated molecules in this system. After infection with the astrocyte network tracer and one week of biotin-supplemented water, mice were perfused with TL. Brains were subsequently stained and cleared. We found no evidence that streptavidin signal followed vasculature in cKO brains (Figure 4k), indicating that vasculature-related mechanisms did not compensate for the loss of astrocyte gap junctions.

### Astrocyte networks are plastic

The barrel cortex is a pliable and accessible forebrain region commonly used to investigate neuroplasticity^28-30^. To test whether astrocyte networks, like neurons, exhibit structural plasticity, whiskers on 4-week-old mice were unilaterally trimmed for 28 days (Figure 5a). This manipulation is known to induce robust structural remodeling in neurons^28-32^. At day 28, a viral mixture containing the astrocyte network tracer and a CaMKIIα-driven mCherry (to label excitatory neurons) was injected into the barrel cortex corresponding to the trimmed whiskers (Figure 5b). Whiskers were trimmed for an additional 4 weeks, with biotin administered via drinking water during the final week.

We quantified network size in both trimmed and naïve conditions by calculating the ratio of streptavidin-positive to HA-positive cells in a cohort with reduced injection volumes to improve measurement precision (Figure 5c and Supplemental Figure 11). Mice with unilaterally trimmed whiskers had significantly smaller astrocyte networks in corresponding barrel cortex (p = 0.002; naïve 3.54 ± 0.39 vs trim 2.16 ± 0.22), indicating that astrocyte networks are indeed structurally plastic.

**Figure 5.**
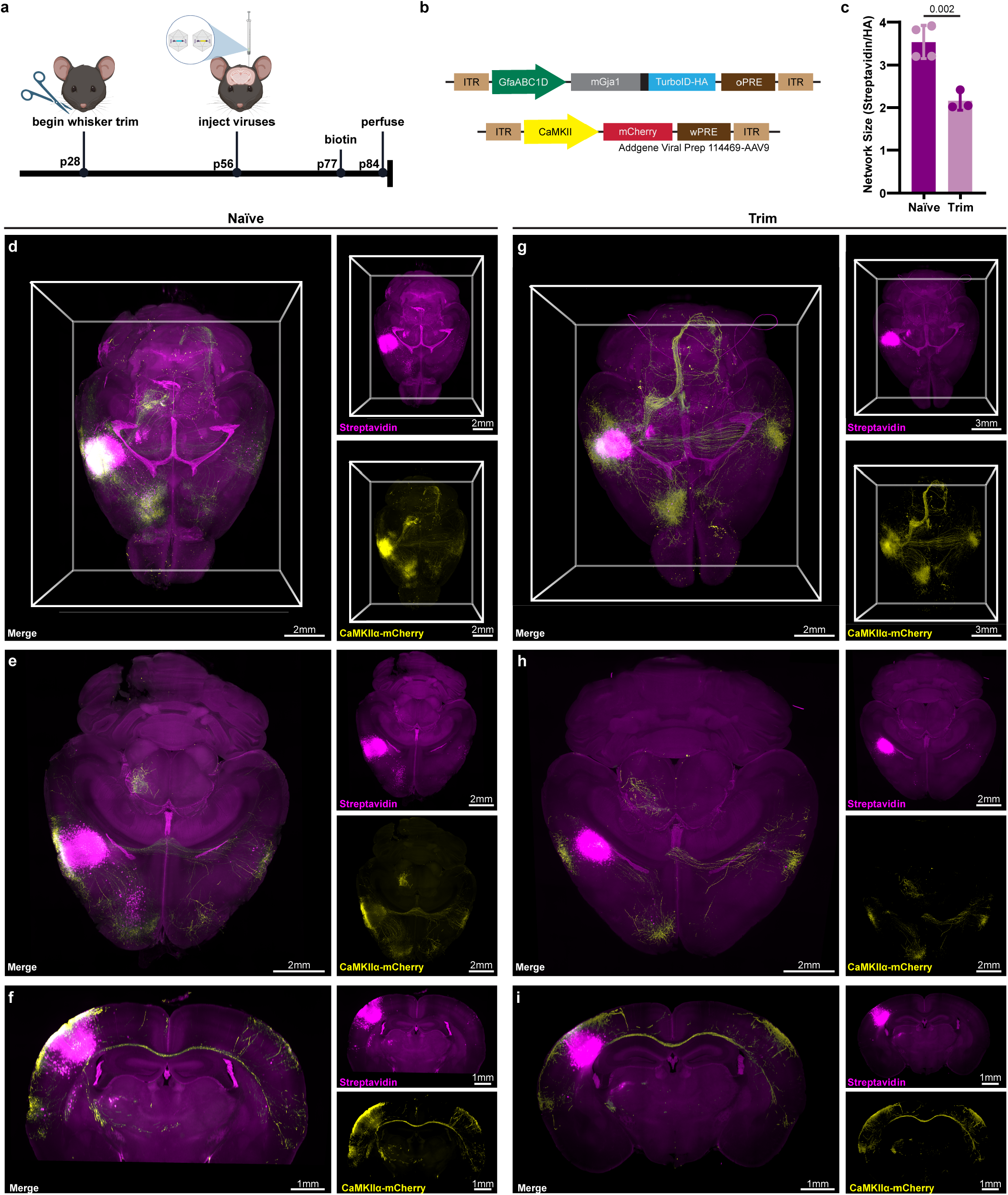
Astrocyte networks are plastic and differ from neuronal networks. **a**. Experimental timeline. **b**. Diagram of the constructs in the viral mixture. **c**. Following whisker trim, the number of streptavidin-positive cells relative to the number of HA-positive cells was significantly reduced in corresponding barrel cortex relative to naïve animals (n = 4 naïve, n = 3 trim) Statistical significance (p value indicated on graph) determined via unpaired Welch ‘s t-test (α = 0.05). Values are mean ± standard deviation. **d**. Three-dimensional rendering of a naïve mouse infected with both constructs in a single injection Streptavidin (magenta) shows astrocyte connectivity originating from the same region in which infected neurons (yellow, mCherry) reside. **e**,**f**. Horizontal (**e**) and coronal (**f**) virtual sections provide a detailed view of connectivity among specific regions. **g-i**. Corresponding images in a brain from a mouse with unilaterally trimmed whiskers corresponding to infected barrel cortex.

### Astrocyte and neuronal networks differ

We next increased the injection volume to assess commonalities and differences among long-range astrocytic networks and neuronal projections. While astrocyte streptavidin signal occasionally overlapped with neuronal projections, this was not uniformly the case. Figure 5d shows a three-dimensional rendering of astrocyte networks and neuronal projections in naïve barrel cortex. While local connections and some long-range projections were shared among astrocytes and neurons, many neuronal projections lacked corresponding astrocyte connectivity. This divergence became more pronounced following whisker trim. As shown in Figure 5g, the astrocyte network contracted markedly, particularly in prefrontal cortex. Virtual horizontal (Figure 5e,h) and coronal (Figure 5f,i) sections of these brains reveal the extent of this effect. In the naïve brain, barrel cortex astrocyte networks project to prefrontal cortex and show limited contralateral signal, both of which are largely absent following whisker trim. In contrast, midbrain connectivity remained relatively stable. These findings demonstrate that long-range astrocyte networks are structurally plastic, and that their architecture can diverge significantly from the neuronal connectome.

## DISCUSSION

Here, we demonstrate that multiple astrocyte networks traverse the mouse brain, each selectively connecting and omitting brain regions rather than diffusing indiscriminately (Figure 2). These networks vary in size and organization, with some local networks limited to a single brain region and other long-range networks robustly interconnecting multiple regions across hemispheres. Interconnected astrocytes display region-specific patterns of connectivity within the same network (Figure 3). In some regions, such as prefrontal cortex in the motor cortex-derived astrocyte network, nearly all astrocytes contain biotinylated molecules. In other regions, only subsets of astrocytes are linked, sometimes in linear chains, other times in honeycomb-like patterns that exclude scattered astrocytes. The morphology of networks in different regions may provide clues about their local functions. In brain regions that are fully interconnected, we hypothesize that the network buffers ions across as many astrocytes as possible. In regions with honeycomb-like patterns, we propose that multiple interleaved astrocyte networks may occupy distinct functional niches for the same neuronal population.

Astrocyte networks can directly link brain regions that are not connected by neurons, suggesting that previously unassociated brain regions communicate with one another through gap junction-coupled astrocytes. What physiological role might such an overlapping multicellular network serve in the brain? The answer to this question is likely multifaceted and beyond the scope of this study. However, it is tantalizing to hypothesize that under physiological conditions, these networks facilitate redistribution of metabolic and antioxidant support from regions of low neuronal activity to those with high demand. Glutathione and phosphocreatine could mediate such support, respectively serving antioxidant and energetic buffering roles and both small enough to diffuse through Cx43 gap junctions^26,33-35^. In contrast, during pathology, astrocyte ynetworks may help dissipate locally accumulated metabolic byproducts or pathogenic molecules by redistributing them across the network for degradation or clearance from the CNS^36^.

Some networks also exhibit streptavidin labeling in small, specific populations of neurons. Connexin 36, the canonical neuronal connexin^37^, is a delta-type connexin and therefore incompatible for heterodimerization with the alpha-type Cx43 or beta-type Cx30 expressed by astrocytes – explaining why neuronal streptavidin staining is so rare. A notable exception is motor neurons, which have been reported to express Cx43 during development or in response to injury^38-40^. Although we don ‘t induce a significant injury and study only adult mice, we observed streptavidin-positive motor neurons projecting through the spinal cord following astrocyte network tracer injection into the motor cortex (Supplemental Figure 8k,n). This observation suggests that connexin-mediated astrocyte-neuron communication in motor neurons may be more prevalent than previously appreciated.

The reason for astrocyte-to-neuron molecular transfer is unclear, but there are several potential explanations. Neurotransmitters have been shown to flux gap junctions^41^, raising the possibility that astrocyte networks resupply neurons with neurotransmitters, allowing them to conserve energy. This may be particularly important for neurons with long, energetically-intensive projections. In addition, highly active neuronal populations may also receive antioxidants such as glutathione (∼307 Da) from astrocytes to mitigate oxidative stress without incurring the metabolic cost of de novo antioxidant production. Future work should use mass spectrometry to identify the specific biotinylated molecules transferred through astrocyte networks, which will help us understand the functional roles of astrocyte networks across conditions.

Crucially for future experimentation, astrocyte networks link both hemispheres, including at least the contralateral counterpart of the infected region. This raises critical considerations: internally controlled experiments must be interpreted cautiously, as astrocyte connectivity may undermine the assumption of regional independence. If a so-called control region is functionally linked to an experimental region by astrocyte networks, it is unlikely to remain biologically naïve.

This bilateral connectivity may also explain the contralateral astrocytic responses to unilateral stressors. For example, ischemic stroke in somatosensory cortex causes astrocytic responses in the contralateral hemisphere following ipsilateral limb stimulation^42^. Similarly, a unilateral model of glaucoma elicits astrocytic responses in both the contralateral retina and the contralateral hemisphere of the brain^11,43^. Importantly, these contralateral responses are not restricted to a single sensory system, suggesting that astrocyte-mediated interhemispheric communication may represent a broader feature of neurodegenerative progression.

We additionally find that astrocyte networks are plastic. Barrel cortex astrocyte networks shrink in response to whisker trim (Figure 5). To mirror the canonical timeline of neuronal remodeling, we trimmed whiskers for a month before astrocyte infection^29^. However, both astrocyte gap junctions and the fine processes that house them can remodel on much shorter timescales. The half-life of Cx43 is estimated to be between 1.5-5 hours^44,45^, and astrocyte process remodeling occurs over similarly rapid intervals^46^. Our whisker barrel experiments indicate that astrocyte networks can reconfigure in a stable manner, but it is also likely that rapid remodeling occurs on a smaller scale during more acute perturbations. Future studies should explore whether acute energetic stress triggers dynamic reorganization of astrocyte networks to meet varied metabolic demands across the CNS.

While our approach visualizes astrocyte networks through a binary lens (connected or not via viral labeling), it is important to recognize that a diverse array of molecules may flux through these networks, and their distribution is unlikely to be uniform. Local and long-distance astrocyte networks may serve distinct physiological roles, even within the same inter-connected chain of cells. Uncovering the specific dynamics of different fluxed molecules and how these dynamics are shaped by the molecular identity and subtypes of each astrocyte will be a complex task. However, such studies may ultimately reveal a rich array of functional niches that astrocyte networks occupy across the brain.

Here, we define the specificity and spatial extent of astrocyte networks across brain regions, demonstrate how they differ from neuronal networks, and reveal their ability to remodel. These findings establish a foundation for future exploration of how astrocyte network structure and function are shaped by injury, disease, development, aging, and experience-dependent processes such as learning and memory.

## METHODS

### Animals

Animal procedures were performed in accordance with National Institutes of Health guidelines with the approval of NYU Grossman School of Medicine ‘s Institutional Animal Care and Use Committee (IACUC). All animals were housed at 22-25 °C with 50-60% humidity. Animals had access to food and water ad libitum and were housed on a 12 hour light/dark cycle. Experiments in Figures 2 and 3 were on male, 12-week-old C57BL/6J mice. For experiments in Figure 4, the Saab laboratory crossbred mice^1^ expressing the tamoxifen-sensitive Cre recombinase Cre-ERT2 under the control of the murine *Slc1a3* (GLAST) promoter^47^ with mice carrying floxed Gjb6 (*Cx30*^fl/fl^)^48^ and floxed Gja1 (*Cx43*^fl/fl^)^49^. Mice hemizygous for *Slc1a3*^Cre-ERT2^ were bred to noncarriers to generate *Slc1a3*^Cre-ERT2+/^*Gja1*^fl/fl^*Gjb6*^fl/fl^ experimental mice and *Gja1*^fl/fl^*Gjb6*^fl/fl^ littermate controls for in vivo experiments; in vivo experiments on this genotype were balanced for sex. When primary mouse astrocytes were isolated, *Slc1a3*^Cre-ERT2^ was kept homozygous to obtain a culture in which all astrocytes could be induced via 4-hydroxytamoxifen. All mice were on a C57BL/6 background. The primer sequences used for genotyping were: for *Cx30* flox; 5 ‘-TTCCCTATGCT-GGTAGAGTGCTTGT-3 ‘and 5 ‘-GCAGTAACTTATTGAAAC-CCTTCACCT-3 ‘; for *Cx43* flox: 5 ‘-GGGATACAGACCCTTG-GACTCC-3 ‘and 5 ‘-TCACCCCAAGCTGACTCAACCG-3 ‘; and for GLAST-CreERT2, 5 ‘-GAGGCACTTGGCTAGGCTCT-GAGGA-3 ‘and 5 ‘-GAGGAGATCCTGACCGATCAGTTGG-3 ‘ and 5 ‘-GGTGTACGGTCAGTAAATTGGACAT-3 ‘. The Saab lab previously analyzed Cre reporter function^1^ by crossing with ROSA26-floxed-STOP-GCaMP6s (Ai96; JAX: 024106). Following shipment, mice were rederived by NYU Grossman School of Medicine ‘s Rodent Genetic Engineering Laboratory.

### Tamoxifen treatment

Tamoxifen treatment was performed as previously described11. For 3 consecutive days, both *Slc1a3*^Cre-ERT2+/^*Gja-1*^fl/fl^*Gjb6*^fl/fl^ experimental and littermate control mice received daily (between 3 and 5 p.m.) gavage of a tamoxifen solution (20 mg/mL dissolved in corn oil). Solution was prepared 2 hours prior to administration, during which it was shaken at 37 °C in the dark. Any further experimental manipulations were performed at least 7 days following the last dose of tamoxifen.

### Primary rat astrocyte cultures

Isolation of primary rat astrocytes by immunopanning was performed as previously described^21^. Briefly, cortices of postnatal day (P) 6/7 Sprague Dawley rats (Charles River) were blunt dissected, meninges removed, and enzymatically dissociated at 37 °C / 10% CO_2_ via papain (Worthington Biochemical LS003126). Tissue was then triturated using a 5mL serological pipette to generate a single cell suspension, resuspended in Dulbecco ‘s Phosphate Buffered Saline (PBS; VWR SH30264.FS) with bovine serum albumin (BSA, Sigma Aldrich A4161) and DNAse (Worthington Biochemical LS002007), and filtered using a 20 μm nitex filter. The suspension was negatively panned for non-specific secondary antibody binding, endothelial cells (BSL-1, Vector Labs L-1100), microglia (CD45, BD Pharmingen 553076), oligodendrocyte lineage cells (O4 hybridoma, generated in house), and positively panned for astrocytes (ITGB5, Thermo Scientific 14-0497-82). Astrocytes were removed from the positive panning plate using TrypLE (Thermo Scientific 12-605-010) and plated at 10K per well in an 8 well chamber slide pre-coated with poly-l-lysine (Ibidi 80804). Astrocytes were cultured in serum-free medium containing 50% neurobasal, 50% DMEM, 100 U/mL penicillin, 100 μg /mL streptomycin, 1 mM sodium pyruvate, 292 μg/ mL L-glutamine, 1× SATO, 5 μg/mL N-acetylcysteine, and 5 ng/ mL HBEGF (Sigma-Aldrich E4643-50UG). After 7 days, AAV was added to the media change, resulting in an effective titer of 105. Cells were incubated for a further 7 days, with biotin (effective concentration 250 μM) added to the media 2 days prior to fixation.

### Expansion microscopy

Expansion microscopy was performed based on published protocols. Primary astrocytes were quickly washed in phosphate buffered saline (PBS), then fixed in 4% PFA in PBS at RT for 15 min. After three 5-min washes in PBS, they were permeabilized for 15 minutes in PBS with 0.5% triton X-100, then blocked for 1 hour in 5% BSA and 0.2% Tween-20 in PBS while shaking at RT. They were then incubated in primary antibody solution overnight at 4°C (3% BSA, 0.2% Tween-20 in PBS containing: Mouse monoclonal anti HA-tag (Cell Signaling 2367) directly conjugated to ATTO488-NHS (ATTO-TEC AD 488-31), Rabbit polyclonal anti-Cx43 (Cell Signaling 3512) directly conjugated to ATTO565-NHS (AT-TO-TEC AD 565-31), and ATTO643-Streptavidin (ATTO-TEC AD 643-61)). Dye conjugation was performed according to the manufacturer ‘s protocol. Antibodies were added for an effective concentration of 2 μM for anti-Cx43, 10 μM for anti-HA, and streptavidin (stock concentration 2ug/ml) was added at 1:1000. Cells were then washed 3 × 5 min in PBS and incubated for 10 min in 300nM DAPI in PBS, then imaged on a spinning disk confocal (CrestOptics X-LIGHT V3 Confocal on Nikon Ti2) with a 60x N.A. 1.4 oil lens to later establish an expansion coefficient. 250 μL Acryloyl-X, SE (Invitrogen A20770) was then added for an overnight incubation at RT. Cells were washed twice for 15 min in PBS, then incubated in 300 μL Gelation Solution per well (For 2 mL: 542 μL 4 M Na Acrylate (VWR S03880), 1mL PROTOGEL (Fisher 50-899-90119), 200 μL 10x PBS, 198 μL H_2_O (MQ), 30 μL 10% ammonium persulfate (Thermo Scientific 17874), 30 μL Tetramethylethylenediamine (TEMED, Thermo Scientific 17919)) for 1 hour at 37 °C. 300 μL digestion buffer was added to each well and gels were carefully outlined with a needle to facilitate detachment; once gels detached, they were each transferred to one well of a 12-well plate for the remainder of the 4 hour incubation at 37 °C (volume per gel/well of 12-well plate: 1550 μL Tris-acetate-EDTA (TAE) buffer, 100μL 10% Triton X-100 in PBS, 320μL 5M NaCl with 26 μL Proteinase K (Thermo Scientific EO0491) added immediately before use). Gels were then each transferred to 15 cm petri dishes filled with RT H2O (MQ) and incubated for 30 minutes at RT. H_2_O (MQ) was then replaced and gels left overnight at RT to expand. Gels were stored at 4 °C until imaging.

### Single-molecule imaging and localization

Gels were mounted into 35 mm dishes with #1.5 coverslip bottoms (Ibidi 81158) with SlowFade Glass soft-set antifade mountant (RI 1.52; Invitrogen S36917) immediately before imaging. For lattice-SIM imaging, we used a ZEISS Elyra 7 with an Alpha Plan Apochromat 63x/1.46 oil TIRF (total internal reflection fluorescence microscopy) objective. Dual PCO.Edge 4.2 sCMOS cameras collected data (1280 × 1280 pixels) with a Dual Camera Beam Splitter (SBS LP 560) utilized for channel separation on Zen Black 3.0 SR software corresponding to 15 phases of a 27.5 μm grating period for both 488 and 561 channels. SIM2 (structured illumination microscopy) 3D Leap processing was run with the following parameters: Input SNR: Low, Iterations: 25, Regularization Weight: 0.01; Processing sampling and output sampling: 2. At least 10,000 frames with a 30ms acquisition time were collected from each sample for each wavelength channel and processed via the Localization Microscopy processing function in 3D. All single- and multi-emitter events were fitted as single emitter events.

### Primary mouse astrocyte cultures

Mouse astrocytes were obtained from *Slc1a3*^Cre-ERT2+/+^ x *Gja1*^fl/fl^*Gjb6*^fl/fl^ (note the homozygous Cre, opposed to hemizygous for in vivo experiments where mice could be genotyped). Cortices of P3 mice were blunt dissected, meninges removed, and enzymatically dissociated at 37 °C / 10% CO_2_ via papain (Worthington Biochemical LS003126). Tissue was then triturated using a 5 mL serological pipette to generate a single cell suspension, resuspended in Dulbecco ‘s PBS (VWR SH30264.FS) with bovine serum albumin (BSA, Sigma Aldrich A4161) and DNAse (Worthington Biochemical LS002007), and filtered using a 20 μm nitex filter. The suspension was negatively panned for non-specific secondary antibody binding, microglia (CD45, BD Pharmingen 553076), oligodendrocyte lineage cells (O4 hybridoma^50^, generated in house), L1 (Millipore Rat anti-L1, MAB5272), and positively panned for astrocytes (HepaCAM51; Human HepaCAM antibody MAB4108, R&D Systems). Astrocytes were removed from the positive panning plate using TrypLE (Thermo Scientific 12-605-010) and plated at 10K per well in an 8 well chamber slide pre-coated with poly-l-lysine (Ibidi 80804). Astrocytes were cultured in serum-free medium containing 50% neurobasal, 50% DMEM, 100 U/mL penicillin, 100 μg / mL streptomycin, 1 mM sodium pyruvate, 292 μg/ mL L-glutamine, 1× SATO, 5 μg/mL N-acetylcysteine, and 5 ng/ mL HBEGF (Sigma-Aldrich E4643-50UG). On day 6, 4-hydroxytamoxifen (Sigma-Aldrich SML1666) was added at 1:5000 (or concentration appropriate for gradient experiments); ethanol:isopropanol (95:5) was added to control wells at the same concentration. On day 7, AAV was added to the media change, resulting in an effective titer of 1.5 × 104. Cells were incubated for a further 7 days, with biotin (effective concentration 250 μM) added to the media 2 days prior to fixation. Cells were washed 2 x in PBS, then fixed in 4% PFA in PBS for 15 minutes prior to immunocytochemistry.

### Immunocytochemistry

Fixed cells were washed 2 x in PBS/azide, then blocked for 2 hours in 5% normal donkey serum (NDS) with 0.1% Tween-20 in PBS/azide. They were then incubated overnight at 4°C with shaking in 3% NDS, 0.1% Triton X-100 in PBS/ azide with corresponding primary antibodies (Rabbit polyclonal Anti-HA: Cell Signaling 3724 at 1:500; Goat polyclonal Anti-GFAP (Glial Fibrillary Acidic Protein): Abcam ab53554 at 1:500). Cells were washed 3 × 5 min in PBS/azide, then incubated for 2 hours at RT with shaking in 1% NDS, 0.1% Triton X-100 in PBS/azide with the corresponding secondary antibodies (Donkey anti-Goat conjugated to Alexa Fluor 488: Invitrogen A-110055 at 1:500; Donkey anti-Rabbit conjugated to Alexa Fluor 555: Invitrogen A-31572 at 1:500) and streptavidin conjugated to AlexaFluor647 (ThermoFisher S32357 at 1:500). Cells were washed 3 × 5 min in PBS/azide, then imaged (2048 × 2048) on a Zeiss 880 laser scanning confocal microscope using a 20x N.A. 0.8 plan apochromat air objective.

### Western Blotting

Cells were lysed in ice-cold 1x RIPA buffer (diluted in ddH2O from 10x, Cell Signaling #9806) with 1 mM PMSF (Cell Signaling #8553) and HaltTM Protease & Phosphatase Single-Use Inhibitor Cocktail (Thermo Scientific #78440). Samples were sonicated and lysate was separated from insoluble material by centrifugation at 20,000 g. 1.5 mm 10% acrylamide gels (Running gel: 4 mL PROTOGEL (Fisher 50-899-90119), 3 mL 1.5 M Tris pH 8.8 + SDS, 5 mL ddH2O, 120 μL 10% ammonium persulfate (Thermo Scientific 17874), and 12 μL Tetramethylethylenediamine (TEMED, Thermo Scientific 17919); Stacking gel: 780 μL PROTOGEL (Fisher 50-899-90119), 1.5 mL 0.5 M Tris pH 6.8 + SDS, 3.75 mL ddH2O, 60 μL 10% ammonium persulfate (Thermo Scientific 17874), and 18 μL Tetramethylethylenediamine (TEMED, Thermo Scientific 17919)). Gels were loaded with 10 μg protein per well and run at 120 V; transfers onto PVDF membranes (Immoblion-FL, IPFL00010; pore size 0.45 μm) occurred overnight at 4°C at 20 V with stirring. Following a 1 hour block (LI-COR Intercept® PBS Blocking Buffer (LI-COR 927-70001)), blots were incubated with rabbit anti-Cx43 (1:1000; Cell Signaling #3512) and mouse anti-Actin (1:1000; abcam ab8226) antibodies in LI-COR Intercept® PBS Blocking Buffer (LI-COR 927-70001) with 0.1% Tween-20 (Sigma #P9416) for 4 hours, then washed 4 × 5 min with PBST. They were then incubated for 1 hour in LI-COR Intercept® PBS Blocking Buffer (LI-COR 927-70001) containing 0.1% Tween20 and 0.1% SDS with donkey anti-mouse IRDye 680LT (1:20,000; LI-COR P/N 926-68072) and donkey anti-rabbit IRDye 800CW (1:20,000; LI-COR P/N 926-32213). Following 3 × 5 min PBST washes and 1 × 5 min PBS wash, blots were imaged on a ChemiDoc MP (BIO-RAD). Band peaks were quantified in FIJI (ImageJ) using the Gels function.

### qPCR

RNA was obtained from cultures using Qiagen ‘s RNeasy Micro Kit (74004) and converted to cDNA via Qiagen ‘s QuantiTect Reverse Transcription Kit (205311) per manufacturer instructions. Real-time PCR was performed using SYBR™ Green Universal Master Mix (ThermoFisher 4309155) on a StepOne Plus Real-Time PCR System (Applied Biosystems) using the following primers: Gja1 (F: TCATTGGGG-GAAAGGCGTGA, R: CATGTCTGGGCACCTCTCTTTCA), *Aldh1l1* (F: TCCCGTCTTTGACCTTGGGT, R: CGCCAC-CGAGGGAACTTAAA), *Slc1a3* (F: CCCCTTACAAAAT-CAGAAAAGTTGT, R: CCCATCTTGGGCTCTTCTCC), *Sox10*: (F: GAAGAAGGCTCCCCCATGTC, R: TTGGGT-GGCAGGTATTGGTC), *Mog* (F: GCAGGTCTCTGTAG-GCCTTG, R: CCCTCAGGAAGTGAGGATCAAA), *Aif1* (F: CTGGGCAAGAGATCTGCCAT, R: ACCAGTTGG-CCTCTTGTGTT), *Cd14* (F: ACTGAAGCCTTTCTCGGAGC, R: TGAAAGCGCTGGACCAATCT), *Tubb3* (F: ACCATGGA-CAGTGTTCGGTC, R: ACACTCTTTCCGCACGACAT), *Syp* (F: CGGCTACCAGCCTGACTATG, R: CTGGGCTTCACT-GACCAGAC).

### Perfusion and tissue preparation

Mice were always perfused between 2 and 5 PM to control for circadian effects. Mice were heavily anesthetized with an overdose of pentobarbital (Euthasol: 390 mg Pentobarbital / 50 mg Phenytoin/mL at 2 uL/g) and transcardially perfused with PBS containing 10 mg/L Heparin (Sigma Aldrich, Heparin sodium salt from porcine intestinal mucosa, H3393) until the liver and right ventricle were clear of blood. Perfusion solution was then switched to 4% PFA in PBS; all solutions were administered at RT at 70% cardiac perfusion pressure. Brains were immediately dissected taking care to leave the surface fully intact and post-fixed in 4% PFA in PBS overnight to stabilize protein tertiary structures. Fixed samples were washed in PBS with 0.01% sodium azide 3 times for 1 hour, then stored in PBS/azide at 4 °C.

When Lycopersicon Esculentum lectin (Tomato lectin, TL) was used to image vasculature, 100 μg of TL conjugated to DyLight 649 diluted in 100 μL PBS was injected directly into the left ventricle of the heart prior to perfusion. After circulating via the beating heart for 1 minute, transcardial perfusion resumed as above.

### Slice Immunohistochemistry

Hemibrains were delipidated (see Clearing), rehydrated through the same methanol gradient to B1n, then washed 3 x in PBS/azide. They were then sectioned sagittally at 200 μm on a vibrating microtome (Leica VT1000 S). Floating sections were blocked for 2 hours in 5% normal donkey serum (NDS) with 0.1% Tween-20 in PBS/azide, then incubated overnight at 4 °C with shaking in 3% NDS, 0.1% Triton X-100 in PBS/azide with corresponding primary antibodies (Rabbit polyclonal Anti-Connexin 43: Cell Signaling 3512 at 1:500; Rabbit polyclonal Anti-Connexin 30: Invitrogen 71-2200 at 1:250; Goat polyclonal Anti-GFAP (Glial Fibrillary Acidic Protein): Abcam ab53554 at 1:500). Sections were washed 5 × 10 min in PBS/azide, then incubated overnight at 4 °C with shaking in 1% NDS, 0.1% Triton X-100 in PBS/azide with the corresponding secondary antibodies (Donkey anti-Goat conjugated to Alexa Fluor 488: Invitrogen A-110055 at 1:500; Donkey anti-Rabbit conjugated to Alexa Fluor 555: Invitrogen A-31572 at 1:500). Sections were washed 5 × 10 min in PBS/ azide and mounted in ProLong Diamond Antifade Mountant with DAPI (Invitrogen P36962) to obtain a close refractive index match to the delipidated sections.

### Confocal Imaging

Sections were imaged on a Zeiss 800 confocal microscope; montages were obtained using a 20x plan apochromat N.A. 0.75 air objective at 512 × 512 per image and high-resolution inset images were obtained using a 63x plan apochromat N.A. 1.4 oil objective at 2048 × 2048 resolution. Z-stacks were obtained at a step size of 8.2 μm (20x) or 4.4 μm (63x). All directly compared sections were obtained at the same magnification and were imaged using the same settings. Maximum Z-projections were rendered using FIJI (ImageJ 1.54f).

### Connexin and vasculature relative localization analysis

Confocal images were analyzed in FIJI/ImageJ (Version 1.54m). Three confocal micrographs for each of three mice per condition were analyzed using ImageJ ‘s ‘Plot Profile ‘ function for three equally spaced line segments perpendicular to imaged vasculature. Line segments were placed so they began on the edge of a vessel, completely transected it, and then passed through the remainder of the micrograph. Profile values for each line were averaged to represent one micrograph. Graph is presented as distance from the far end of each blood vessel with the average in-vessel portion of all images demarcated.

### Intracerebral Viral Injections

Surgeries were performed under aseptic conditions in accordance with NYU Grossman School of Medicine ‘s institutional biosafety guidelines. A glass micropipette was pulled to a tip diameter of approximately 20 microns, filled with mineral oil, and attached to a Nanoject III microinjector (Drummond Scientific, 3-000-207). The micropipette was then backfilled with 1ul of Adeno-associated Virus (AAV) at a titer of 1010 genomic units per microliter. Mice were anaesthetized with 0.7-2.5% isoflurane (adjusted on the basis of scored reflexes and breathing rate during surgery), placed into a stereotaxic apparatus (Kopf), and a single craniotomy performed over the experimental brain region (coordinates below), the micropipette was inserted, and 200 nL of virus was injected (40 cycles, 5 second delay between cycles, 5 nL injected per cycle and 5nL/s). 10 minutes following injection the micropipette was withdrawn and the wound sutured closed. Buprenorphine (0.1 mg/kg) was subcutaneously administered immediately following surgery and twice daily for the following 72 hours. We used the following stereotaxic coordinates (in mm: AP, anteroposterior; ML, mediolateral; DV, dorsoventral): Motor Cortex (AP -2.6; ML 1.3; DV -0.8), Hippocampus (AP -2.0; ML 1.5; DV -1.5), Paraventricular Nucleus (AP 0.12; ML -0.71, DV -4.8), Prefrontal Cortex (AP 0.35; ML 1.9; DV -2.5), Barrel Cortex (AP -1.5; ML 3.5; DV -0.7). The astrocyte network tracer (AAV5-GfaABC1D-Cx43:TID:HA) was constructed and packaged by VectorBuilder. The AAV9-CaMKIIa-mCherry was obtained from Addgene as a gift from Karl Deisseroth (Addgene viral prep #114469-AAV9). All vectors were injected at a titer of 1 × 211.

### Biotin administration

Biotin (Sigma Aldrich, B4639) stocks were diluted to 100 mM in DMSO, aliquoted for single-use, and stored frozen at -20 °C. 21 days following intracerebral injections, biotin stock was diluted in distilled water to 160 μM (400 μL of 100 mM stock per 250 mL of distilled water) and provided ad libitum for 7 days, after which mice were perfused (always between 2 and 5 p.m. to control for circadian effects).

### Whisker trimming

Every other day from P28 through P84, mice were lightly anaesthetized with 0.7-1.5% isoflurane (adjusted on the basis of scored reflexes and breathing rate). All whiskers one side of the face were trimmed with scissors. On P56, whisker trim coincided with intracerebral viral injections (see above). For P77-P84, mice received biotin-supplemented drinking water (see above); perfusion occurred on P84 (see above).

### Clearing

The clearing protocol uses similar principles to the iDISCO^10^ or AdipoClear^9^ protocol with several adjustments specific to brain tissue and astrocyte morphological maintenance. To delipidate, fixed samples were washed in B1N buffer (H2O / 0.1% Triton X-100 / 0.3 M glycine / 0.01% sodium azide, pH 7) for 1 hour, followed by 1 hour washes each in 20%, 40%, 60%, and 80% methanol in B1N. Samples were fully dehydrated in 100% methanol (MeOH) for three 1 hour washes, then delipdated in 2:1 dichloromethane (DCM; Sigma-Aldrich SHBR8133) overnight. Delipidation resumed the following day in three 1 hour washes in 100% DCM. Samples were then washed in 100% MeOH for 1 hour twice. Residual fluorescence was quenched through an overnight incubation in 80% MeOH / 5% H_2_O_2_ / 15% H_2_O at 4 °C. Brains were then rehydrated via 1 hour incubations in each 60%, 40%, and 20% methanol in B1N. Samples were then washed in B1N overnight.

Samples were permeabilized in PTxwH buffer (PBS / 0.1% Triton X-100 / 0.05% Tween-20 / 2 μg/mL heparin / 0.02% sodium azide) with an added 45 g/L glycine and 50 mL/L dimethyl sulfoxide (DMSO) two times for 1 hour each. Then, samples were washed three times in PTxwH for 1 hour.

Samples were incubated in primary antibody (HA-tag, Cell Signaling mAb #3724, 1:2000) and conjugated streptavidin (Alexa Fluor 647 Streptavidin, ThermoFisher S32357, 1:800) for 1 week at 37 °C with rotation. After primary incubation, samples were washed in PTxwH with 8 solution changes over a period of 3 days. Samples were then incubated in secondary antibody (donkey anti-rabbit conjugated to AlexaFluor Plus 555, ThermoFisher A32794) and conjugated streptavidin (1:800) for 1 week at 37 °C with rotation. As after secondary incubation, samples were washed in PTxwH with 8 solution changes over a period of 3 days.

For tissue clearing, samples were dehydrated in a 20%, 40%, 60%, 80%, 100%, 100%, 100% methanol/H2O series for 45 minutes each. Following dehydration, samples were washed in 2:1 DCM:methanol overnight followed by three 1 hour washes in 100% DCM. Samples were cleared overnight in dibenzyl ether (DBE, Sigma-Aldrich) and stored at room temperature in the dark in amber glass vials until imaging.

### Light sheet imaging

Whole, cleared brains were imaged on a light sheet microscope (Zeiss Z1) equipped with a 5x EC plan neofluar N.A. 0.16 objective lens and 20x Clr plan neofluar N.A. 1.0 objective lens. Images were collected with two 1920 × 1920 pixels sCMOS cameras and acquired using Zen Black software (Zeiss). Samples were staged in a custom imaging chamber (Translucence Biosystems) filled with DBE and illuminated from both sides by the laser light sheet (5x light sheet thickness:13.08 μm, step size 4.5 μm; 20x light sheet thickness: 4.35 μm, step size 0.61 μm).

### Light sheet image analysis

For cKO experiments, Imaris (Version 9.9) was used to generate surfaces around positive signal (calculated via thresholding) in HA and Streptavidin channels. The total volume of Streptavidin signal divided by the total volume of HA signal was used to generate a Relative Network Volume representing each brain. For whisker trim experiments, individual astrocytes were counted in FIJI (ImageJ version 1.54b) following background thresholding. The number of streptavidin positive astrocytes was divided by the number of HA positive astrocytes to generate a Network Size value representing each brain.

### Statistical analysis and reproducibility

Data are presented as mean ± standard deviation. Mice were assigned at random to groups. Experiments were not performed in a blinded fashion. Statistical significance for each experiment was determined as detailed in the respective figure legend. In figure legends, n = the biologically independent number of animals per group. Statistical analyses were performed in Prism 10 software (GraphPad). No data points were excluded.

## Acknowledgements

We thank the Microscopy Core at New York University (NYU) Langone Medical Center for experimental and technical support. We thank Giorgia Bertoli for her advice on expansion microscopy and the course directors of the Kavli iDISCO course at Rockefeller University for their expertise in tissue clearing. We thank the NYU Langone Medical Center Genotyping Core and Mouse Genetic Engineering Core for their assistance with colony rederivation and genetic maintenance. We thank Eleni Katafygiotou, Brianna Somoza Balza, Jesus Morales Santos, and Adwoa Amponsah for their assistance with tissue clearing and Jessica Minder for her cloning and surgical expertise. We thank Simon Peron for his advice on working with whisker barrel cortex. We thank Paul Glimcher, Stephen Davis Jr, and Xavier Castellanos for their proofreading assistance. Thanks go to the Lean Levy Fellowship in Neuroscience (M.L.C.), the PEW Charitable Trusts Postdoctoral Fellowship (M.C.S.), the Simons Foundation SURFiN Fellowship (K.E.C./M.V.C.), the Swiss National Science Foundation (320030-232028, A.S.S.) and the NIH [K00 AG068343 (H.K.G.), 5U19 NS107616 (M.V.C.), T32 MH019524 (M.L.C.), and R01 EY033353 (S.A.L.)]. NYU shared resources are partially supported by the Cancer Center Support Grant P30CA016087 at the Laura and Isaac Perlmutter Cancer Center. Thanks additionally go to the Belfer Neurodegeneration Consortium of MD Anderson (S.A.L.), the Carol and Gene Ludwig Family Foundation (S.A.L.), and the Cure Alzheimer ‘s Fund (S.A.L.).

## Author Contributions

M.L.C. designed research; S.A.L. and M.V.C. supervised the work; M.L.C., M.C.S., M.C., H.K.G., J.S., and K.E.C. performed research; M.L.C. and A.S.S. contributed new reagents/analytic tools; M.L.C. analyzed and visualized the data; M.L.C. wrote the paper; M.L.C., M.C.S., M.C., H.K.G., K.E.C., A.S.S., S.A.L., and M.V.C. edited the paper; M.L.C., M.C.S., H.K.G., S.A.L., and M.V.C. acquired funding.

---

**Supplemental Figure 1.**
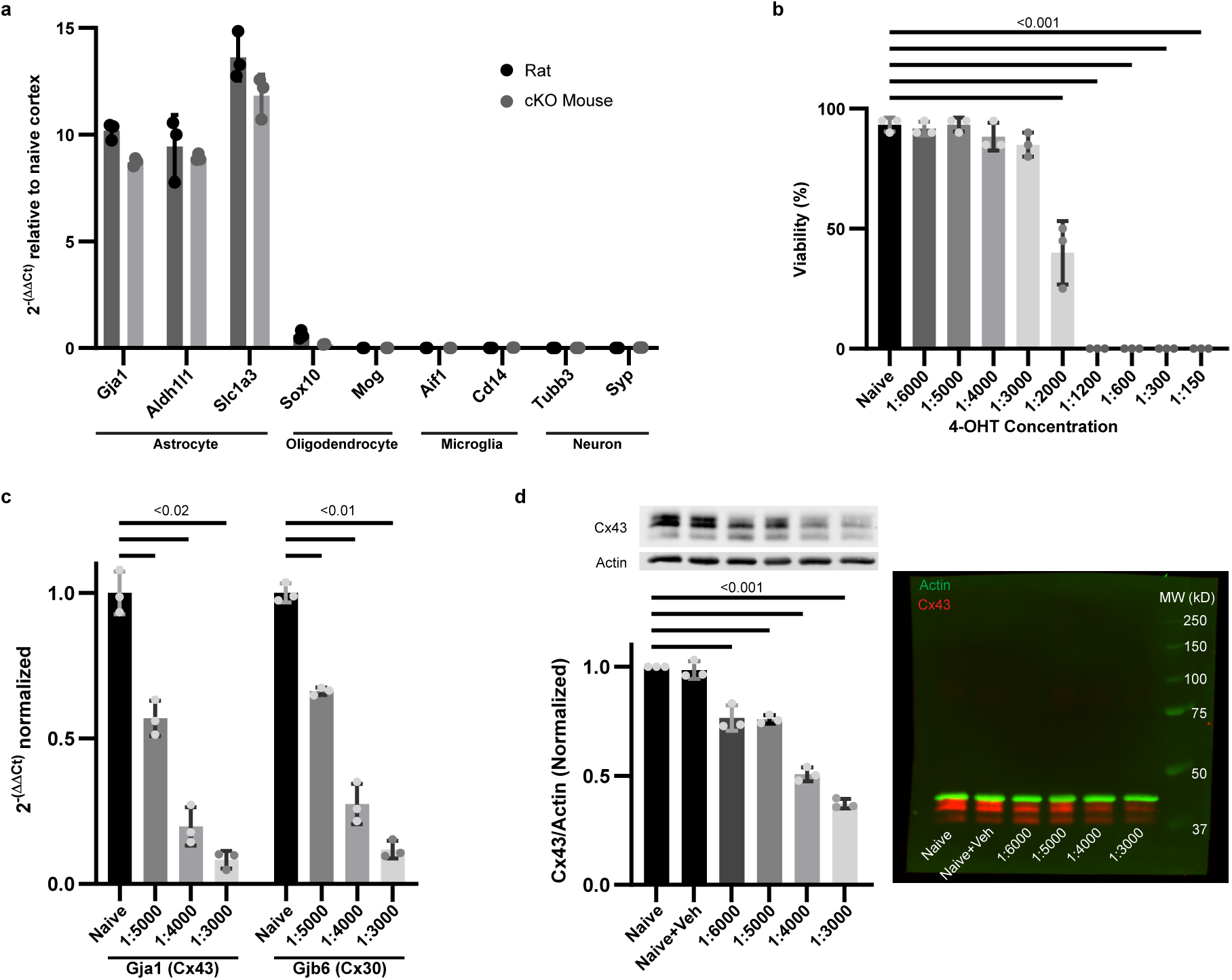
Primary immunopanned astrocyte purity and knockout induction. **a**. qPCR of marker gene expression for astrocytes (*Gja1, Aldh1l1, Slc1a3*), oligodendrocytes (*Sox10, Mog*), microglia (*Aif1, Cd14*), and neurons (*Tubb3, Syp*) in primary immunopanned astrocytes from rat or cKO mouse normalized to the ribosomal gene RPL19 and relative to naïve cortex from simultaneously dissected littermate controls. **b**. Primary cKO mouse astrocyte viability determined as the ratio of Hoechst positive, Propidium Iodine (PI) negative cells to the total number of Hoechst positive cells. Viability was significantly decreased at 4-Hydroxytamoxifen (4-OHT) concentrations of 1:2000 and above (p < 0.001). **c**. qPCR of *Gja1*(Cx43) and *Gjb6* (Cx30) in primary immunopanned cKO astrocytes at a 4-OHT concentrations of 1:5000, 1:4000, and 1:3000 relative to naïve cells. All 4-OHT concentrations decreased expression of both RNAs, with the largest decrease at 1:3000. d. Quantification of western blots from immunopanned cKO astrocytes in the following conditions: Naïve, Naïve + Vehicle (EtOH, 1:3000), 4-OHT 1:6000, 4-OHT 1:5000, 4-OHT 1:4000, and 4-OHT 1:3000. Blots were costained for Cx43 and Actin (example on right). Cx43/Actin normalized to Naïve significantly decreased at all 4-OHT concentrations, with the largest decrease at 1:3000. n = 3 independent isolations for all panels. Statistical significance was determined by repeated measures one-way ANOVAs (p values indicated on graphs) followed by Dunnett ‘s multiple comparisons test (α = 0.05). Values are mean ± standard deviation.

**Supplemental Figure 2.**
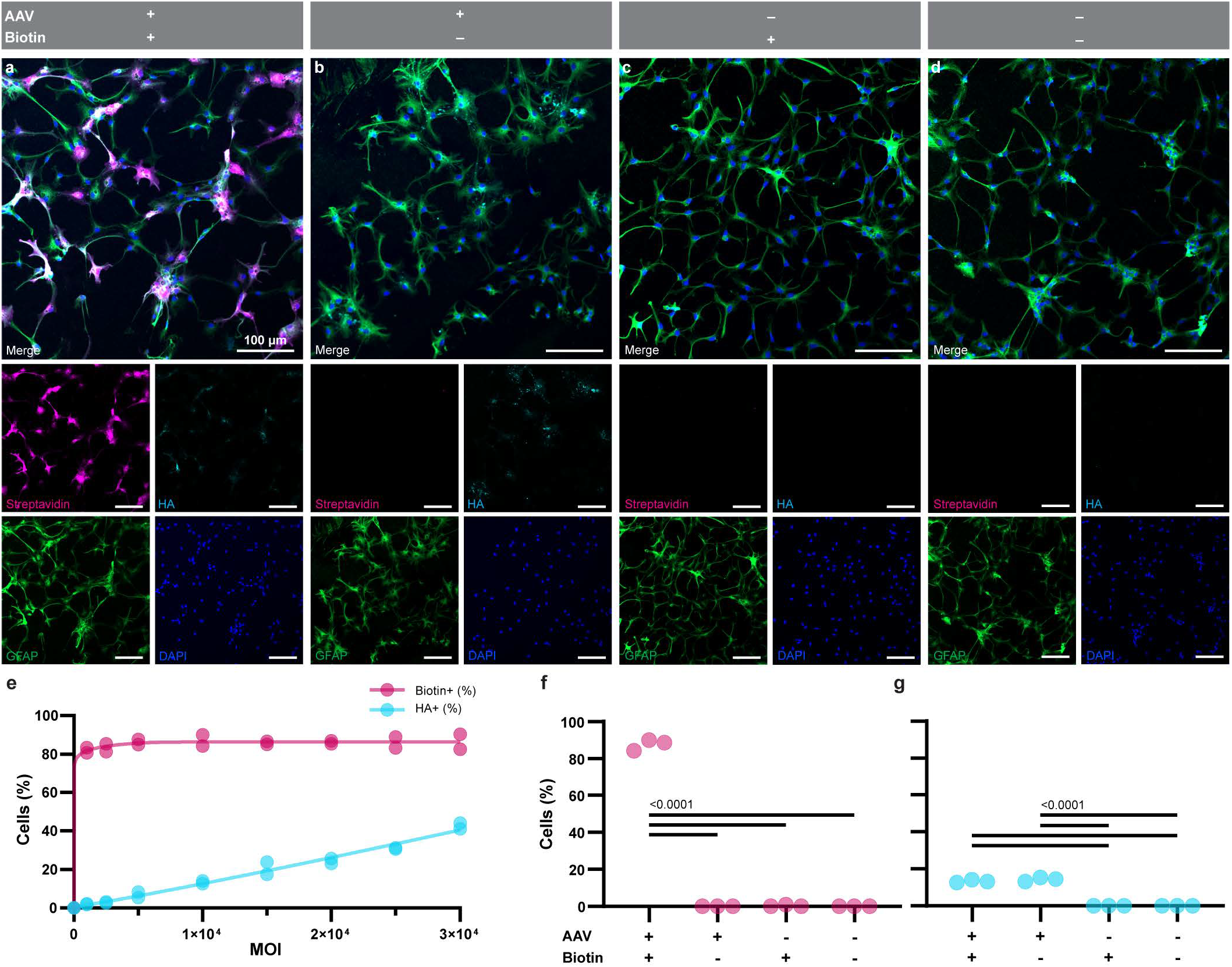
Both AAV5-GfaABC1D-Cx43:TID:HA and biotin are required for astrocyte network detection. **a**. (also in Figure 1e) Primary rat astrocytes (GFAP, green) infected with AAV5-GfaABC1D-Cx43:TID:HA and given biotin have punctate staining patterns for HA-tag in about 20% of cells; ∼80% of cells are positive for streptavidin. **b**. Without supplemented biotin, HA-tag is detected but not streptavidin. Without AAV (**c**) or with neither AAV nor biotin (**d**), neither HA-tag nor streptavidin are detected. **e**. Primary rat astrocytes were incubated with a range of MOIs (multiplicity of infection) to estimate the viral load necessary for ∼10% of astrocytes to express Cx43-TID. Across each MOI tested, about 80∼ of astrocytes were positive for biotinylated fluxed molecules. n = 2 isolations f. 87.7 ± 3.0% of cells from the AAV+/Biotin+ condition (MOI = 1 × 104) were positive for streptavidin; the other 3 conditions contained no detectable streptavidin signal. g. 13.3 ± 0.8% of cells were positive for HA-tag in both AAV+ conditions; neither AAV-condition contained detectable HA-tag immunopositivity. n = 3 isolations for both g and h. Statistical significance (p values for all comparisons < 0.0001) determined via one-way ANOVA followed by Tukey ‘s multiple comparisons test (α = 0.05).

**Supplemental Figure 3.**
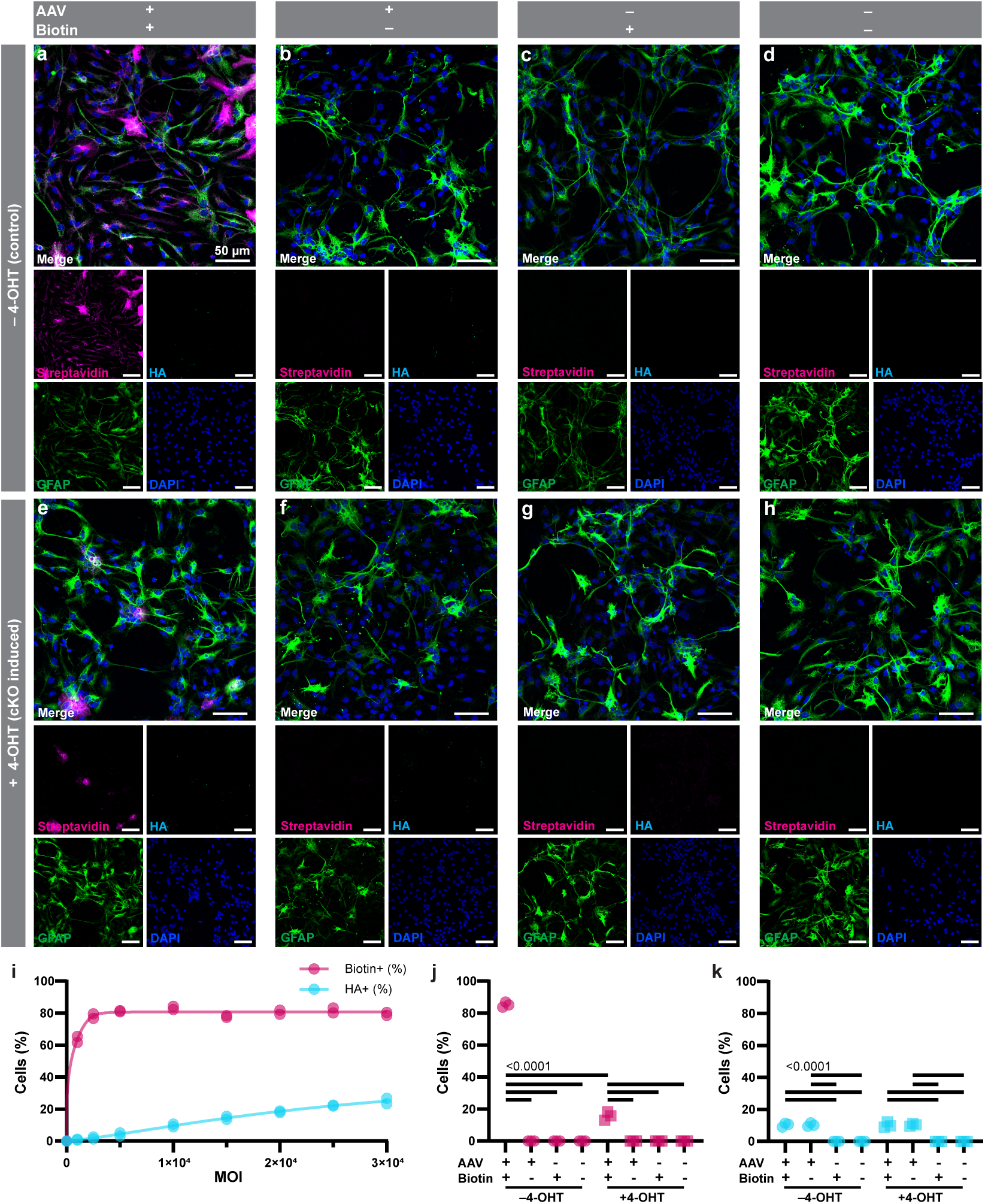
Genetically-expressed Cx43 is necessary for biotin to distribute beyond infected cells. **a**. Primary astrocytes isolated from *Slc1a3*^Cre-ERT2+/+^*Gja1*^fl/fl^*Gjb6*^fl/fl^ mice (GFAP, green) demonstrate distributed streptavidin staining and punctate HA-tag staining when infected with AAV5-GfaABC1D-Cx43:TID:HA and given biotin. **b**. Only HA-tag is visible when AAV is given without biotin. Without AAV (**c**) or with neither AAV nor biotin (**d**), neither HA-tag nor streptavidin are detected. **e**. Astrocytes induced for (*Gja1*) and Cx30 (*Gja6*) knockout via 4-OHT are only positive for streptavidin in HA-tag positive cells in the AAV and biotin condition. As in b-d, only HA-tag is visible when AAV is given without biotin (**f**). Without AAV (**g**) or with neither AAV nor biotin (**h**), neither HA-tag nor streptavidin are detected. **i**. Primary mouse astrocytes isolated from uninduced cKO mice were incubated with a range of MOIs to estimate the viral load necessary for ∼10% of astrocytes to express Cx43-TID. Across each MOI tested other than 1 × 103, about 80% of astrocytes were positive for biotinylated fluxed molecules; for 1 × 103, ∼60% of astrocytes were streptavidin positive. **j**. 85.2 ± 1.6% of cells from the control AAV+ and Biotin+ condition (MOI = 1 × 104) were positive for streptavidin; the other 3 control conditions contained no detectable streptavidin signal. For cKO astrocytes induced with 4-OHT, 15.7 ± 2.6% of cells from the AAV+ and Biotin+ condition were positive for streptavidin; the other 3 cKO condition contained no detectable streptavidin signal. k. ∼10% of cells were positive for HA-tag in all AAV+ conditions (control++ 10.3 ± 1.2%; control+-10.4 ± 1.1%; cKO++ 10.3 ± 1.7%; cKO+-10.4 ± 0.8%); no AAV-condition contained detectable HA-tag immunopositivity. n = 3 isolations for both j and **k**. Statistical significance (p values for all comparisons <0.0001) determined via one-way ANOVA followed by Tukey ‘s multiple comparisons test (α = 0.05). Values are mean ± standard deviation.

**Supplemental Figure 4.**
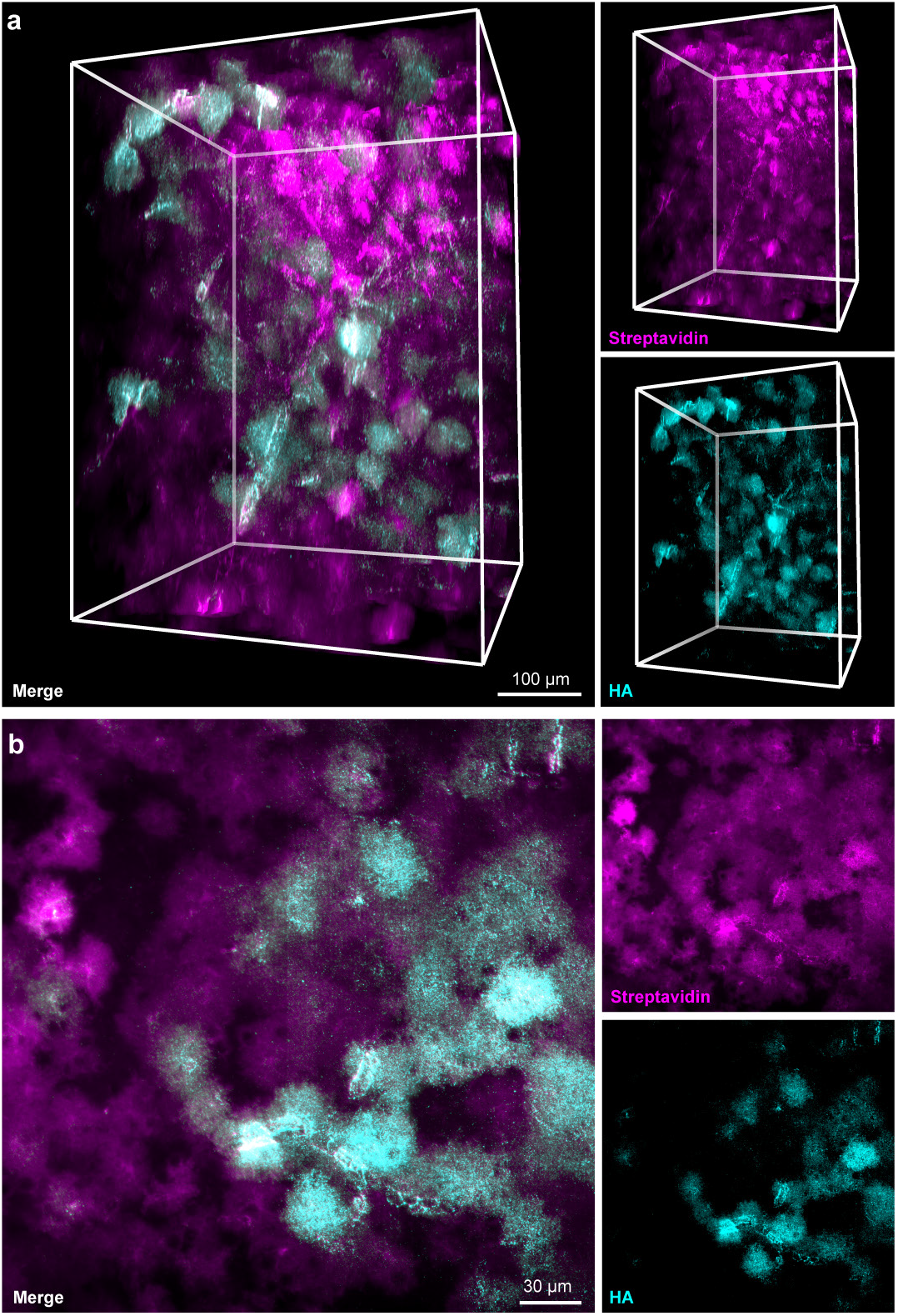
Infected cells exhibit astrocytic morphology and punctate HA-tag staining. **a**. Three-dimensional rendering of infected cells in motor cortex stained for streptavidin (magenta) and HA-tag (cyan). **b**. Single slice shows neighboring infected and uninfected cells. All cells demonstrate astrocytic morphology when filled with biotinylated molecules (streptavidin, magenta); infected cells exhibit a punctate pattern of HA-tag staining (cyan) on the surface of each cell indicative of gap junctional localization.

**Supplemental Figure 5.**
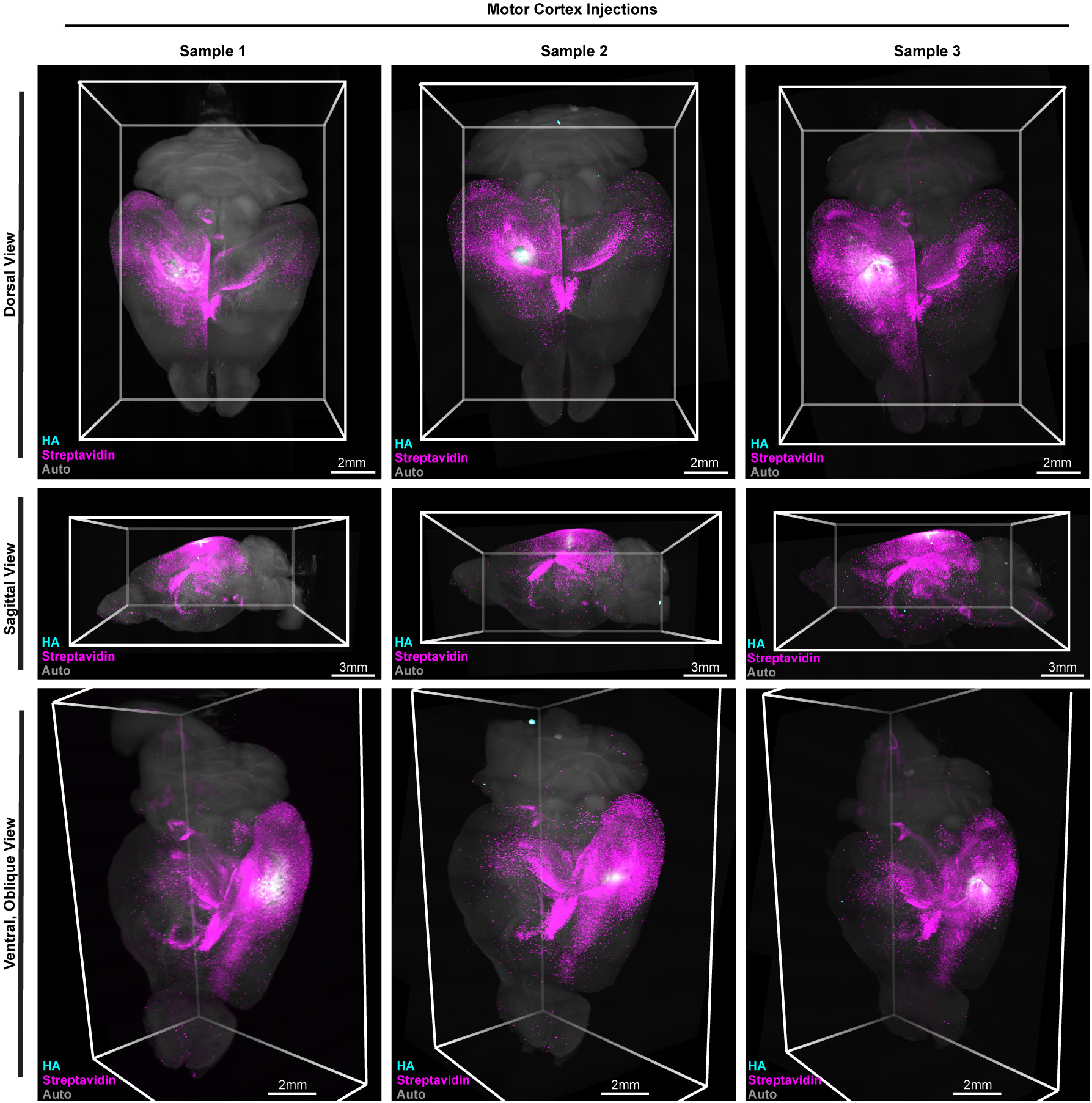
Motor cortex network reproducibility. Light sheet imaged brains from 3 independent mice (columns) shown in dorsal (top row), sagittal (middle row), and ventral/oblique views (bottom row). The infection site is stained for HA-tag (cyan), the astrocyte network originating in the infection site is stained with streptavidin (magenta), and an empty channel ‘s autofluorescence (grey) shows the outline of the brain.

**Supplemental Figure 6.**
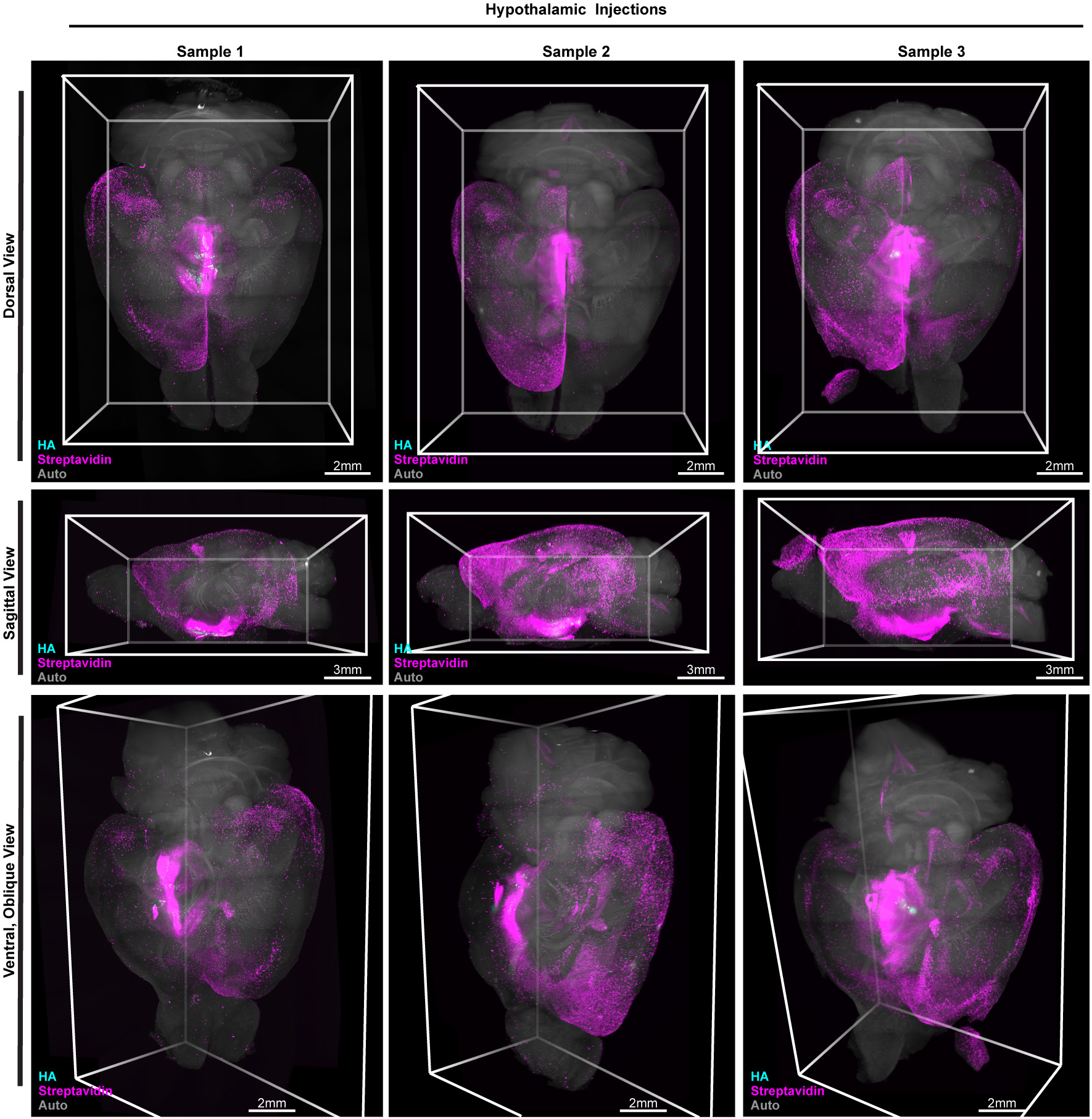
Hypothalamic network reproducibility. Light sheet imaged brains from 3 independent mice (columns) shown in dorsal (top row), sagittal (middle row), and ventral/oblique views (bottom row). The infection site is stained for HA-tag (cyan), the astrocyte network originating in the infection site is stained with streptavidin (magenta), and an empty channel ‘s autofluorescence (grey) shows the outline of the brain.

**Supplemental Figure 7.**
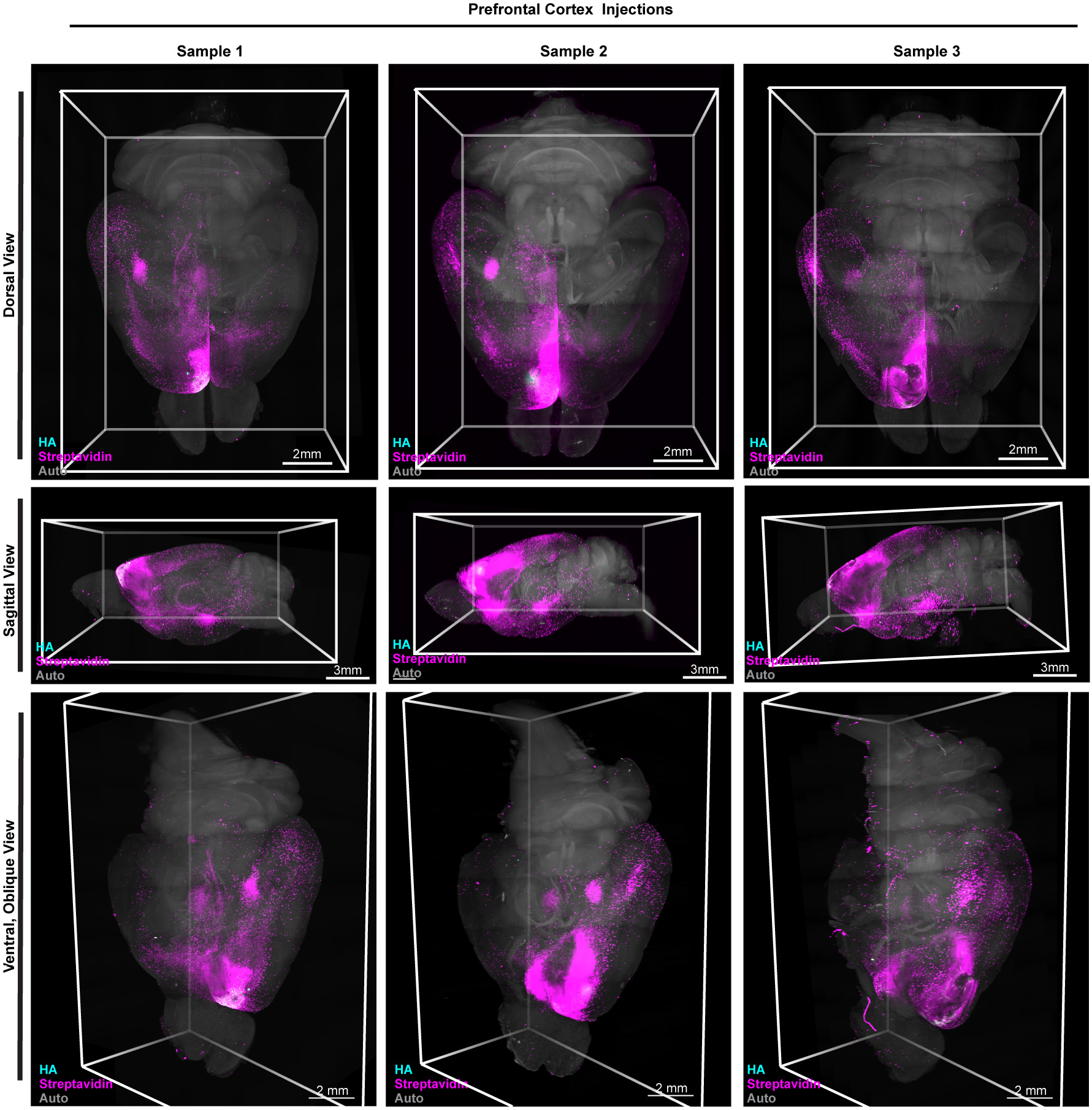
Prefrontal cortex network reproducibility. Light sheet imaged brains from 3 independent mice (columns) shown in dorsal (top row), sagittal (middle row), and ventral/oblique views (bottom row). The infection site is stained for HA-tag (cyan), the astrocyte network originating in the infection site is stained with streptavidin (magenta), and an empty channel ‘s autofluorescence (grey) shows the outline of the brain.

**Supplemental Figure 8.**
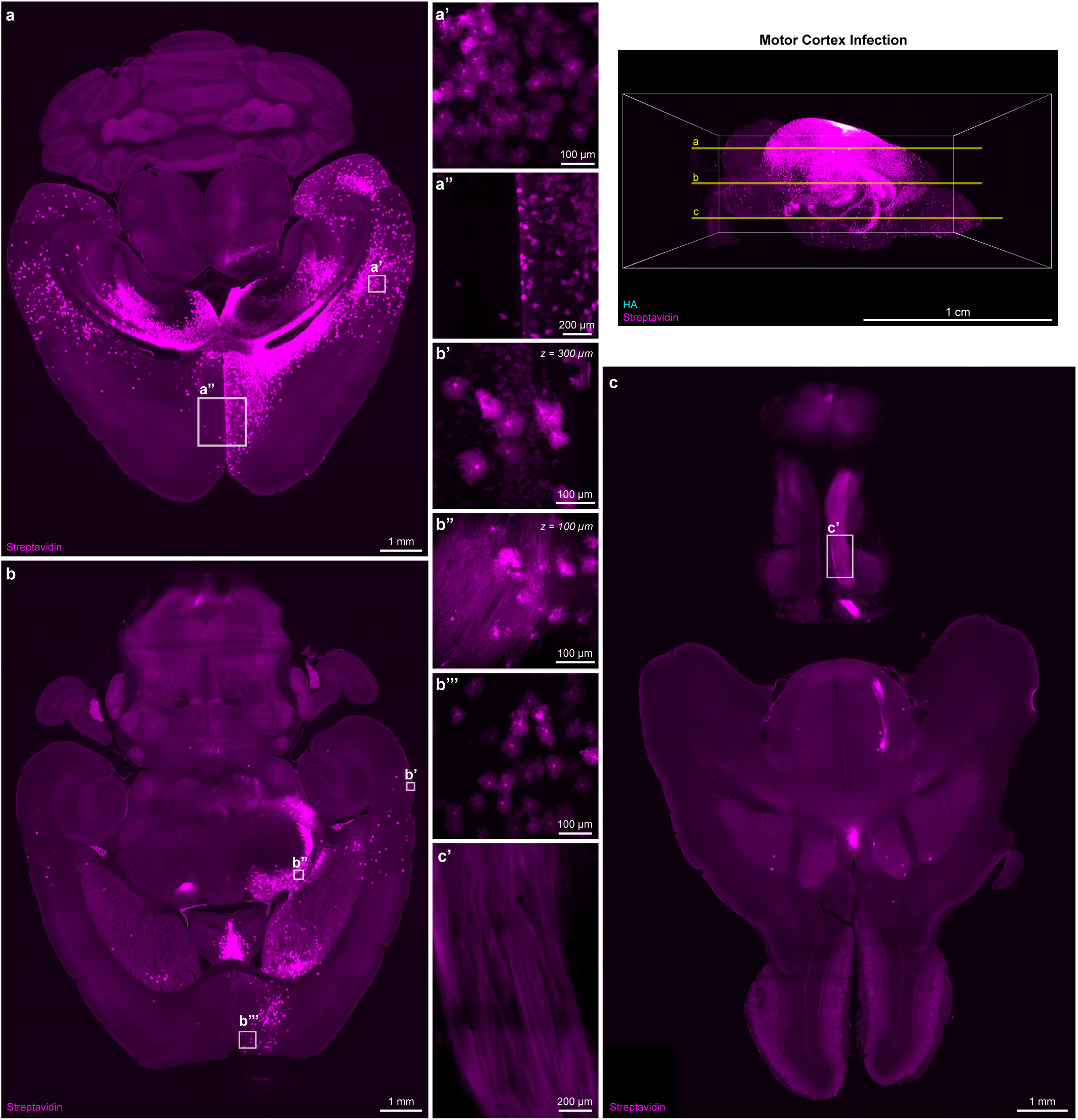
Diversity of cell types within a motor cortex astrocyte network. **a, b, c**. Individual slices at varied depths in the horizontal plane through a light sheet imaged brain infected in motor cortex (yellow lines in reference 3D image, top right; HA, cyan; Streptavidin, magenta). **a ‘**. Biotinylated compounds fill many, but not all, adjacent cells. **a”**. While many brain regions show signal bilaterally, some regions, such as prefrontal cortex in this brain, do not. **b ‘**. Following motor cortex infection, sparse fluorescence in some neurons is visible. **b”**. Chains of astrocytes contact and diverge from neuronal axons, which qualitatively become brighter following astrocyte network contact points. **b ‘ ‘ ‘**. Most astrocyte network termini, such as this terminus in prefrontal cortex, do not show detectable streptavidin signal among adjacent neurons. **c ‘**. Following motor cortex infection, axons descending to the spinal cord show visible streptavidin signal.

**Supplemental Figure 9.**
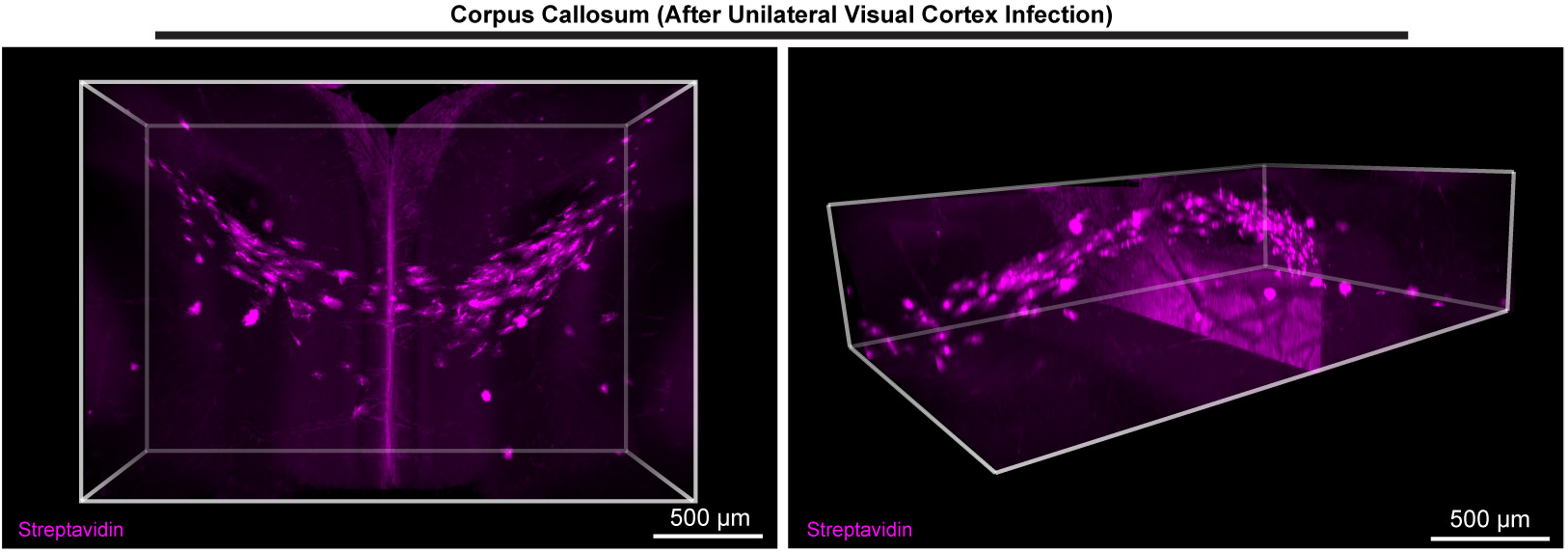
Chains of fibrous astrocytes interconnect bilateral astrocyte networks. Three-dimensional rendering (horizontal and oblique) of streptavidin-positive astrocytes in corpus callosum following infection in visual cortex.

**Supplemental Figure 10.**
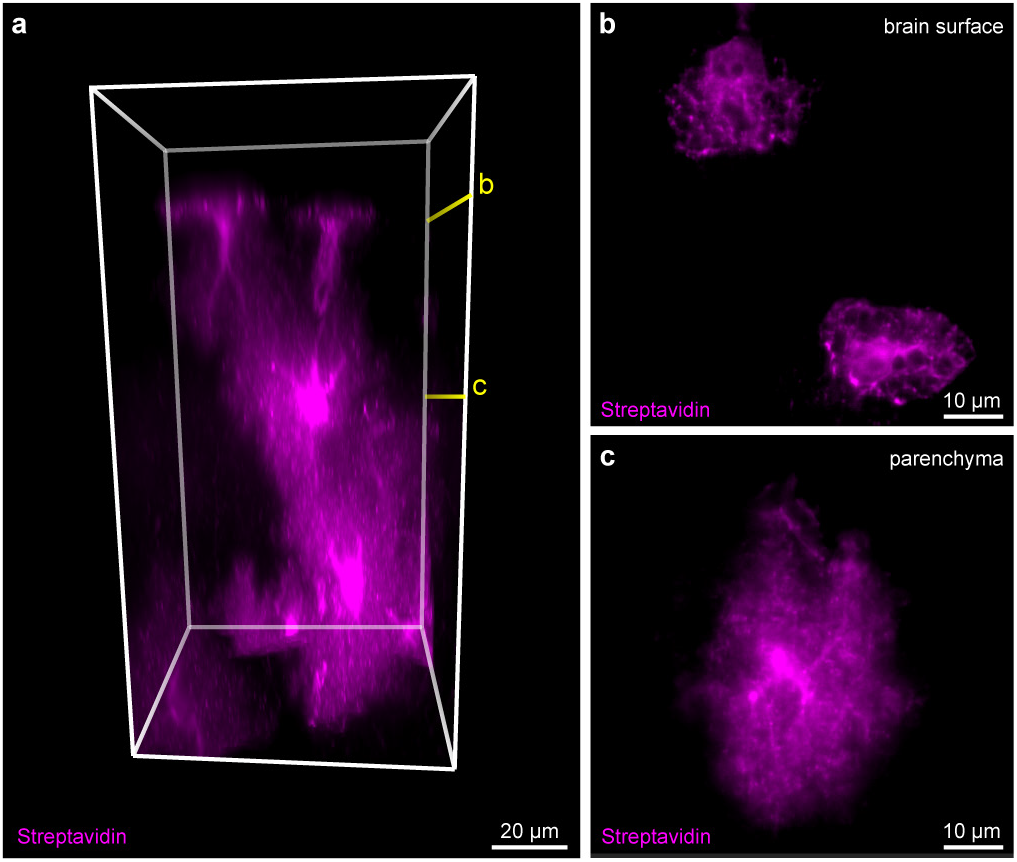
Cortical astrocyte networks can contain both parenchymal and glia limitans superficialis astrocytes on the brain ‘s surface. **a**. Three-dimensional rendering of streptavidin positive astrocytes on the surface of the brain (top) connected to astrocytes in the parenchyma. **b**. Slice (marked in **a**) showing glia limitans superficialis astrocytes on the brain ‘s surface. **c**. Slice (marked in **a**) showing the parenchymal astrocyte in contact with the glia limitans superficialis astrocytes.

**Supplemental Figure 11.**
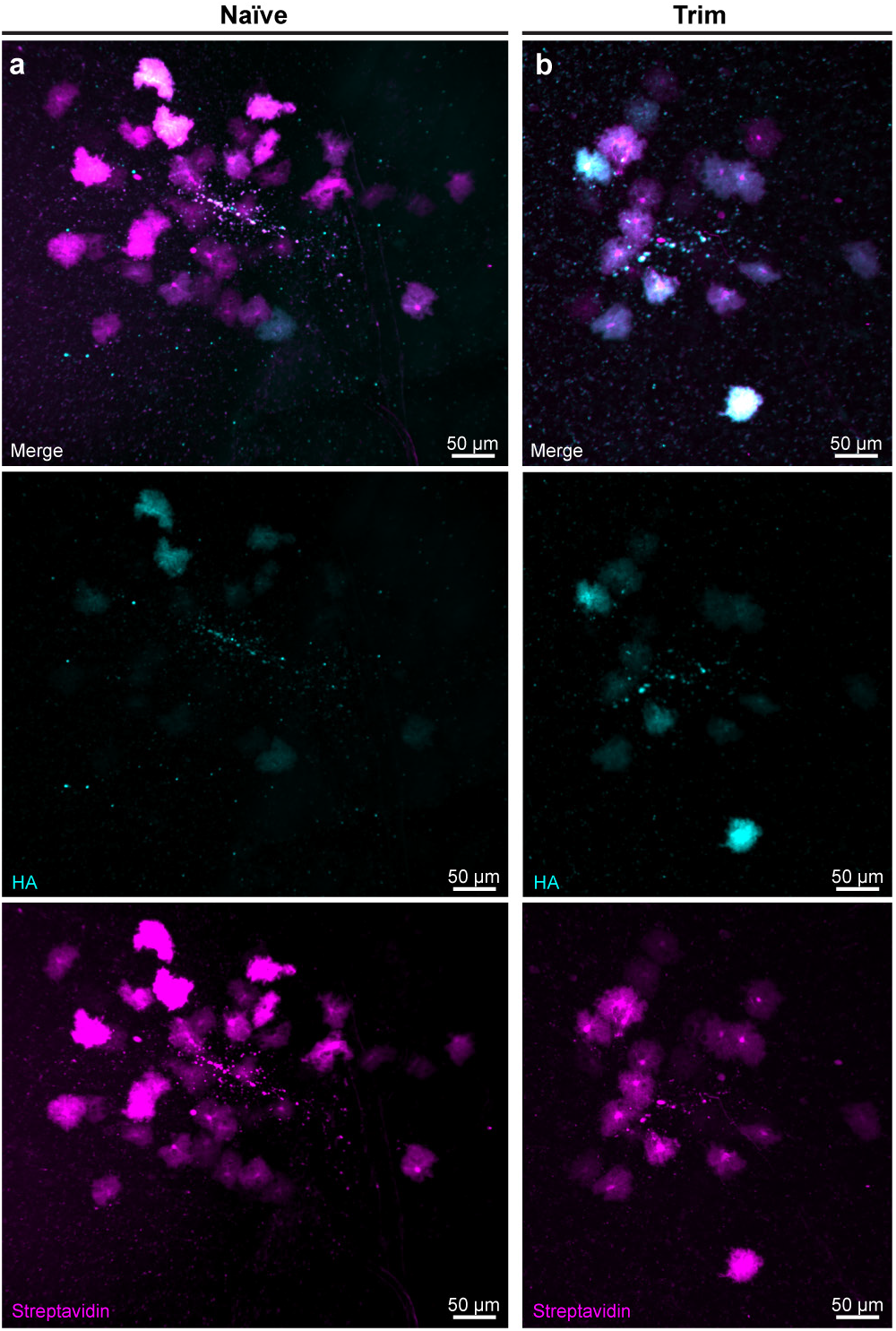
Whisker trim reduces astrocyte network size. Z-projections of all fluorescent cells in animals infected to express *Cx43*-TID-HA (cyan) in barrel cortex, resulting in biotinylation of fluxed molecules in an astrocyte network (magenta). Small-volume (20 nl) injections into barrel cortex corresponding to naïve (**a**) or trimmed whiskers (**b**) resulted in cells labeled at a sparse enough density to be counted and quantified (see Figure 5c).

